# Immunoinformatics-Guided Design and In Silico Evaluation of a Multi-Epitope Vaccine Against Influenza A H10N5 and H3N2 Strains Based on Hemagglutinin and Neuraminidase Proteins

**DOI:** 10.64898/2026.07.03.736294

**Authors:** Muhammad Zeeshan Shabbir, Prem Kumar, Muhammad Atiq Ur Rehman, Jeevan Kumar, Umama Urooj, Syeda Izza Batool, Chandra Sourav, Rumesa Ghazanfar, Zainab Nagari, Daniyal Hameed, Abdul Wahid, Ayesha, Muhammad Daniyal Siddique

## Abstract

Influenza A viruses H3N2 and H10N5 represent, respectively, a persistently dominant seasonal pathogen and a newly documented zoonotic threat with the latter strain variants responsible for the first confirmed human fatality in January 2024, yet no vaccine platform currently addresses co-protection against both subtypes within a unified immunogen. We report here the immunoinformatics based vaccine design and multi-layered computational validation of a 419-amino-acid multi-epitope subunit vaccine construct targeting conserved hemagglutinin (HA) and neuraminidase (NA) antigens identified through multiple sequence alignment of the avian H10N5 (A/swine/Hubei/10/2008) and H3N2 human reference strain sequences to identify viral agents undergoing mammalian adaptations. Linear B-cell, cytotoxic T lymphocyte (CTL), and helper T lymphocyte (HTL) epitopes were predicted using ABCpred, BCEpred, BepiPred 2.0, NetMHCpan 2.1, and NetMHCpan 4.0, then filtered through VaxiJen 3.0, AllerTOP v2.1, and ToxinPred to retain only antigenic, non-allergenic, non-toxic candidates. The final construct, incorporating an avian β-defensin N-terminal adjuvant with GPGPG, AAY, and EAAAK linkers, exhibited a molecular weight of 43.9 kDa, instability index of 31.15, and SOLPro solubility probability of 0.763. Tertiary structure modeling via I-TASSER and GalaxyRefine achieved 84.4% Ramachandran-favored residues. Molecular docking against TLR3 and TLR7 yielded binding free energies of −16.1 and −16.8 kcal/mol with picomolar dissociation constants. Molecular dynamics simulations confirmed complex stability over extended trajectories. Furthermore, codon optimization produced a Codon Adaptation Index of 1.0 for E. coli K12 expression. In silico immune simulation demonstrated robust activation of humoral and cellular immunity including elevated IgG1, IgM, IFN-γ, IL-2, rapid NK cell expansion, and broad B-cell clonal diversity. These findings establish a computationally validated candidate capable of providing protection against influenza in multiple host organisms, warranting experimental advancement.

## Introduction

Influenza (flu) is one of the most persistent viral infections affecting humans, with a history of recurring seasonal outbreaks and occasional pandemics (Javanian, Barary et al. 2021). Influenza viruses belong to the Orthomyxoviridae family and possess segmented, negative-sense, single-stranded RNA genomes comprising approximately 13.5 kb nucleotides. Influenza viruses are classified into three types (A, B, and C) based on differences in their nucleoprotein (NP) and matrix (M) proteins (Dunning, Thwaites et al. 2020). Influenza A and B viruses are responsible for seasonal human epidemics, whereas influenza C typically causes mild seasonal infections (Petrova and Russell 2018). The virus presents a significant global health challenge due to its capacity for rapid mutations and efficient transmission. According to the World Health Organization (WHO), the annual burden of influenza is substantial, with 3 to 5 million severe cases and up to 650,000 influenza-related respiratory deaths (Fischer, Gong et al. 2014). Despite advances in surveillance, vaccines, and antiviral therapies, influenza continues to cause widespread illness and mortality (Meseko, Sanicas et al. 2023). Seasonal mortality patterns are influenced by environmental factors such as humidity and temperature variations, as well as sudden changes in viral virulence resulting from genetic mutations or reassortment events (Neumann and Kawaoka 2022). The virus also possesses a substantial ability to infect a variety of organisms including aves (Williams, Sánchez-Llatas et al. 2023).

Infection typically causes sudden onset of fever, cough, runny nose, fatigue, headache, and sore throat, usually after an incubation period of 1 to 3 days (El-Radhi 2019). Severe infections may progress to life-threatening complications, including neurological aberrations, respiratory distress syndrome, and sepsis, ultimately resulting in death (Garg, Jain et al. 2025).

Recent reports document the circulation of multiple influenza strains, including the human seasonal H3N2 strain and emerging avian-related subtypes such as H10N5 (Houta, Ahmed et al. 2025). The H3N2 strain has been one of the dominant seasonal human influenza viruses in recent years, frequently associated with severe outbreaks, particularly in vulnerable populations including the elderly and young children (Jester, Uyeki et al. 2020, Scarvaglieri, De Francesco et al. 2026). The H10N5 subtype, though less common in human populations, has raised significant concerns due to sporadic detections in birds and the potential for cross-species transmission to humans (Hao, Wu et al. 2025).

Recently, the avian specific H10N5 strain has emerged as a potential threat to human health. A severe case of avian H10N5 infection was documented in a 60-year-old woman in Anhui Province, China, in January 2024, representing the first reported human case and resulting in a fatal outcome. The H10N5 virus predominantly binds avian-type receptors and lacks known mammalian-adapted mutations (Yang, Zheng et al. 2025). Phylodynamic analysis revealed that the H10N5 virus emerged in late 2022 as a result of multiple reassortment events involving strains circulating in Japan, South Korea, Central Asia, and China (Yuan, Zhang et al. 2024). The PA gene was acquired from H5N2 viruses in domestic waterfowl, while seven gene segments were obtained from H10N7 or other low-pathogenic avian influenza viruses carried by wild waterfowl (Boonyapisitsopa, Chaiyawong et al. 2024). The HA gene of the H10N5 virus belongs to the North American lineage, which was likely introduced into Asia by migratory birds and subsequently established local circulation (Wu, Zhang et al. 2025).

Given the widespread impact of H3N2 and the emerging threats of H10N5 subtypes demonstrating mammalian adaptations, which can lead to more severe disease outcomes, effective vaccination strategies are essential to prevent co-infection and reduce associated complications (Nafiz, Marlia et al. 2024). Currently, therapeutic options for influenza rely primarily on vaccination, which remains the most effective preventive measure (Nypaver, Dehlinger et al. 2021). Recent studies demonstrate that seasonal influenza vaccines generally reduce the risk of illness or hospitalization by 40% to 60% (Nichol and Treanor 2006, Monto 2010). Seasonal vaccines are available in two forms: trivalent formulations containing two influenza A strains (H1N1 and H3N2) and one influenza B lineage, and quadrivalent formulations that include both B lineages for broader protection (Tsilibary, Charonis et al. 2021). However, these vaccines have significant limitations (Rolfes, Flannery et al. 2019, Fisman, Pérez-Rubio et al. 2025). Frequent genetic changes in influenza viruses, particularly antigenic drift, can create mismatches between vaccine composition and circulating strains, reducing protection in some seasons (Kim, Webster et al. 2018, Petrova and Russell 2018). Additionally, the immunity provided is relatively short-lived, typically lasting less than one year, necessitating annual re-immunization (Chan, Alizadeh et al. 2021). Immune responses vary considerably among populations, with young children, older adults, and immunocompromised individuals demonstrating reduced vaccine effectiveness (Kunisaki and Janoff 2009). Furthermore, inequitable access in many low-and middle-income countries limits the global impact of vaccination efforts (Kraigsley, Moore et al. 2021). These limitations underscore the urgent need for improved next-generation vaccines with broader and more durable protection (Ashraf, Raza et al. 2025).

Peptide-based vaccines for influenza show considerable promise for targeting key viral proteins and providing precise, broad protection (Herrera-Rodriguez, Meijerhof et al. 2018, Malonis, Lai et al. 2019, Jahantigh, Rezanavaz Gheshlagh et al. 2026). For H3N2 and H10N5 co-infection prevention, peptide-based vaccine development has gained attention for generating focused immune responses (Wang, Tan et al. 2010, Nafiz, Marlia et al. 2024). Several studies have employed immunoinformatics techniques to design multi-epitope vaccines against viral and bacterial infections, including influenza (Maleki, Russo et al. 2021, Sharma, Kumari et al. 2021, Sethi, Varghese et al. 2024). Currently, no human transmission of H5N10 avian strain (A/swine/Hubei/10/2008) has been reported. However, due to its prevalence in aves and swine, the viral agent needs continuous monitoring.

In this study, we developed an in silico multi-epitope vaccine targeting the human H3N2 and avian H10N5 subtype (A/swine/Hubei/10/2008) co-infection. The notion was to capture viruses which demonstrated mammalian adaptation and to address the critical need for developing effective influenza vaccines capable of preventing infection in diverse hosts while making use of the recent advancements in immunoinformatics based vaccine designing (Ghazanfar, Nagari et al. 2026). Through detailed immunoinformatics analysis, we identified two essential viral proteins, hemagglutinin (HA) and neuraminidase (NA), as primary vaccine targets due to their vital roles in viral entry and release, respectively, as well as their strong antigenic potential (Wohlbold and Krammer 2014, Creytens, Pascha et al. 2021).

## Material and Methods

### Protein Sequence Retrieval

The amino acid sequences for neuraminidase (NA) and hemagglutinin (HA) proteins of the avian influenza H10N5 reference strain (A/swine/Hubei/10/2008) were retrieved from the National Center for Biotechnology Information (NCBI) using the accession numbers AFR42736.1 (472 amino acids) and AFR42734.1 (561 amino acids) respectively. Multiple sequence alignment was performed using Clustal Omega to identify conserved regions between the NA and HA proteins of H10N5 reference strain and their homologous proteins in Influenza A virus avian flu-related H3N2 strain (NCBI accession no. HA: JX308801.1, NA: JX286593.1) and human H3N2 strain proteins (See Table 1) (Sievers and Higgins 2018). Based on the conservation analysis, the conserved regions in both HA and NA genes were subjected to epitope predictions.

**Table 1.**
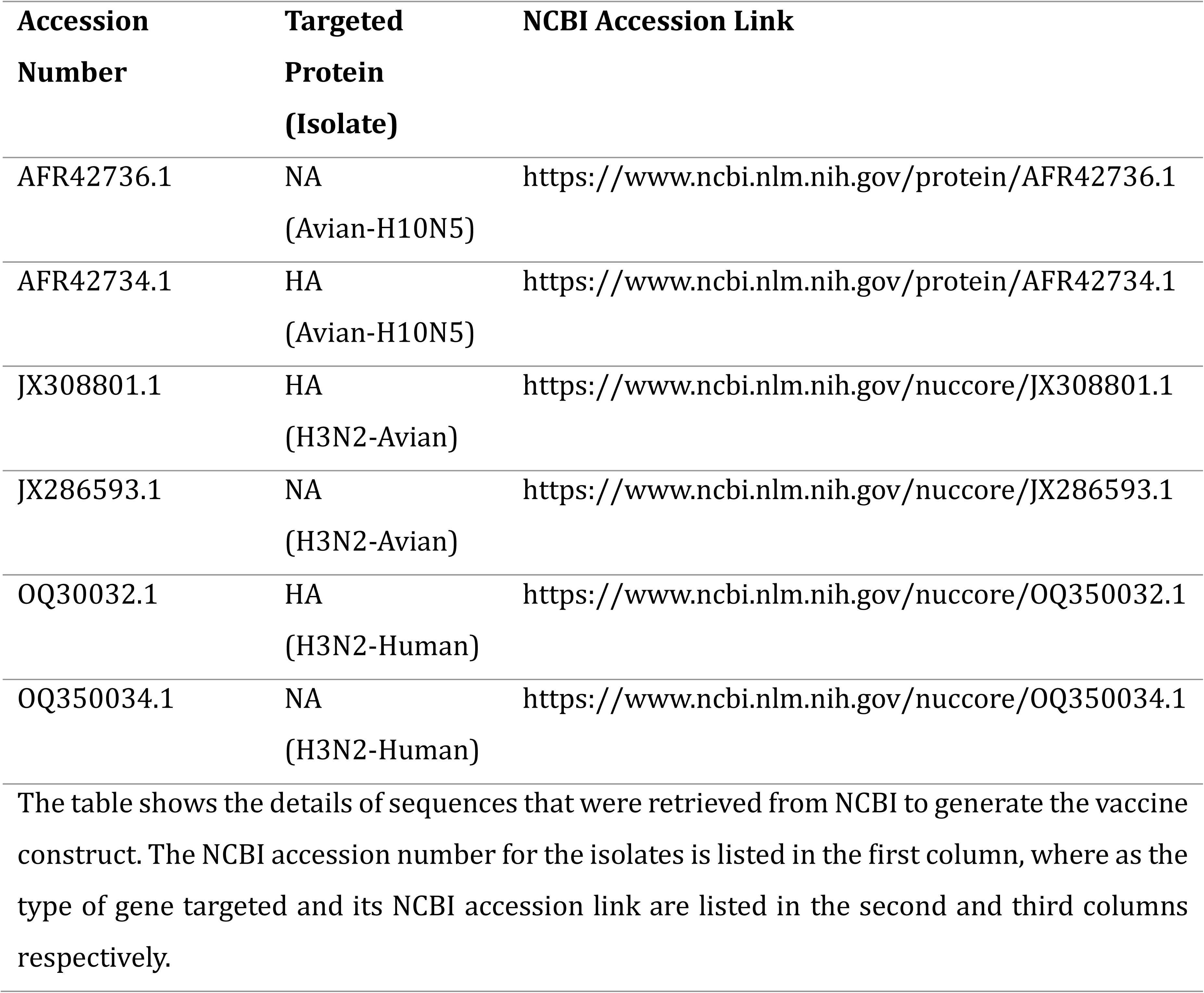
NCBI GenBank accession details for the sequences used for vaccine construct generation.

| **Accession Number** | **Targeted Protein (Isolate)** | **NCBI Accession Link** |
| --- | --- | --- |
| AFR42736.1 | NA<br>(Avian-H10N5) | https://www.ncbi.nlm.nih.gov/protein/AFR42736.1 |
| AFR42734.1 | HA<br>(Avian-H10N5) | https://www.ncbi.nlm.nih.gov/protein/AFR42734.1 |
| JX308801.1 | HA<br>(H3N2-Avian) | https://www.ncbi.nlm.nih.gov/nuccore/JX308801.1 |
| JX286593.1 | NA<br>(H3N2-Avian) | https://www.ncbi.nlm.nih.gov/nuccore/JX286593.1 |
| OQ30032.1 | HA<br>(H3N2-Human) | https://www.ncbi.nlm.nih.gov/nuccore/OQ350032.1 |
| OQ350034.1 | NA<br>(H3N2-Human) | https://www.ncbi.nlm.nih.gov/nuccore/OQ350034.1 |

The table shows the details of sequences that were retrieved from NCBI to generate the vaccine construct. The NCBI accession number for the isolates is listed in the first column, where as the type of gene targeted and its NCBI accession link are listed in the second and third columns respectively.

### B-cell Epitope Prediction

A comprehensive computational pipeline was established to identify and validate linear B-cell epitopes. Initially, ABCpred was used to predict linear B-cell epitopes using artificial neural network algorithms (Saha and Raghava 2006). The prediction window was set to 20 amino acids, and peptides with scores ≥ 0.5 were considered to have high immunogenic potential and were advanced for further analysis. Also BepiPred 2.0, which integrates hidden Markov models with propensity scales, was employed to cross-validate epitopes identified by ABCpred (Jespersen, Peters et al. 2017). This approach provided probabilistic scoring for each residue, enabling the selection of regions with consistent epitope potential. BCEpred was used as an additional validation layer (Saha and Raghava 2004). This tool predicts linear B-cell epitopes using physicochemical parameters including hydrophilicity, flexibility, surface accessibility, polarity, and antigenic propensity. Only regions that demonstrated consistently high scores across multiple parameters were retained for further consideration.

### CTL Epitopes

Cytotoxic T lymphocyte (CTL) epitopes are short peptide regions of proteins recognized by CD8+ T cells (cytotoxic T cells), triggering immune responses that identify and eliminate infected cells. To facilitate vaccine development, probable CTL epitopes were predicted for NA and HA proteins using NetMHCpan 2.1 server (Trolle, Metushi et al. 2015), with default parameters and a sequence length of 15 amino acids. NetMHCpan 2.1 uses neural network-based models to estimate peptide binding affinities to MHC class I molecules across a wide range of HLA alleles. The predicted epitopes were further filtered using VexiJen 3.0 (Doytchinova and Flower 2007), AllerTOP v2.1 (Dimitrov, Bangov et al. 2014), and ToxinPred (Sharma, Naorem et al. 2022) to assess immunogenicity, allergenicity, and toxicity, respectively.

### HTL Epitopes

Helper T lymphocyte (HTL) epitopes are short peptides presented by MHC class II molecules that stimulate CD4+ T helper cells, which coordinate various arms of the immune system. HTL epitopes were predicted using NetMHCpan 4 with default parameters and a sequence length of 12 amino acids, considering antigenicity score, binding score, and IC50 value as key parameters. NetMHCpan 4.0 (Yang, Wei et al. 2024) predicts MHC class II-restricted peptides across HLA-DR,-DP, and-DQ alleles with high precision by integrating binding affinity and eluted ligand data.

### Selection of Epitopes

The antigenicity of all predicted epitopes was assessed using the VaxiJen platform. A threshold of ≥ 0.5 was applied to select peptides likely to be antigenic based on their physicochemical profiles. To eliminate epitopes with potential toxicity, ToxinPred was employed. This tool uses support vector machine (SVM)-based algorithms to identify toxicity based on peptide composition. Only non-toxic peptides were retained for downstream analysis. Furthermore, allergenicity assessment was performed using AllerTOP v2.1. Epitopes predicted as non-allergenic were deemed acceptable for inclusion in the vaccine construct.

### Vaccine Construct Design

To maintain a suitable vaccine size along with assurance of antigenicity, non-toxicity and non-allergenicity of the vaccine construct, only specific epitopes were incorporated into the final vaccine construct based on their antigenicity scores. Strategic linkers were employed to connect the vaccine components: the GPGPG linker was used for epitope-to-epitope linkage of both B-cell and CTL epitopes, while the AAY linker was used for HTL epitopes. These linkers serve dual functions of preventing epitope fusion and facilitating proper antigen processing. To assemble the final construct, the EAAAK linker was used to sequentially join the epitope clusters, with B-cell epitopes linked to CTL epitopes, and CTL epitopes linked to HTL epitopes, ensuring structural continuity and stability within the construct. The N-terminal region of the vaccine was linked to the β-defensin avian adjuvant through an EAAAK linker (Soman, Nair et al. 2009), while the C-terminal region was connected to a 6×His Tag for subsequent purification using the same linker (Janknecht, de Martynoff et al. 1991).

### Physicochemical Properties

Physicochemical properties of the vaccine construct were estimated using the ExPASy ProtParam online server (Gasteiger, Hoogland et al. 2005). Parameters, including the molecular formulae, molecular weight, isoelectric point, instability index (II), number of atoms, estimated half-life and grand average of hydropathicity (GRAVY), were estimated. Also, the SolPro server was used to access the solubility of the vaccine construct to ensure its efficient expression in living cells (Magnan, Randall et al. 2009).

### Secondary Structure Analysis

Self-optimized prediction method (SOPMA) with default parameters was used to predict the secondary structure of the vaccine construct (Geourjon and Deleage 1995). Estimations for proportions of alpha helix, beta sheets, beta-turns and random coils were made. Additionally, PSIPRED provided insights into the structural elements and amino acid composition of the vaccine (McGuffin, Bryson et al. 2000).

### Tertiary Structure Prediction

The tertiary structure of the vaccine construct was predicted using the I-TASSER server (Iterative Threading Assembly Refinement; https://zhanggroup.org/I-TASSER/) (Zhang 2008). Five structural models were generated upon completion of the analysis. These models were subsequently evaluated using the PROCHECK server to assess stereochemical properties, and the top-ranked model was selected for further refinement (Laskowski, MacArthur et al. 1993). The selected model was refined using the GalaxyRefine tool, which generated five refined structural models (Heo, Park et al. 2013). The highest quality refined model was further validated using the ProSA-web server for structural quality evaluation and the PROCHECK server for stereochemical assessment (Wiederstein and Sippl 2007). The final refined structure was visualized using ChimeraX to examine surface topology, backbone geometry, and potential active sites (Pettersen, Goddard et al. 2004).

### Molecular Docking

Molecular docking was performed between the proposed vaccine construct and Toll-like receptors (TLRs) particularly TLR3 and TLR7 respectively. The three-dimensional structures of the TLR3 (PDB ID: 2A0Z) and TLR7 (PDB ID: 5GMH) were retrieved from the Protein Data Bank (PDB). The vaccine construct model and the TLR structures were uploaded to the ClusPro 2.0 web server, an automated, widely-used protein–protein docking platform (Kozakov, Hall et al. 2017). Following docking, the resulting vaccine-TLR3 and vaccine-TLR7 complexes were analyzed for binding affinity and molecular interactions using the PRODIGY and PDBsum servers (Xue, Rodrigues et al. 2016, Laskowski, Jabłońska et al. 2018).

### Molecular Dynamics Simulation

Molecular dynamics simulations were conducted to evaluate the stability and conformational dynamics of the vaccine-TLR3 and vaccine-TLR7 complexes over extended time scales. The top-ranked docked complexes from the ClusPro 2.0 analysis were subjected to molecular dynamics simulations using Schrödinger (Kosloff and Kosloff 1983). Simulations were performed in explicit solvent at physiological temperature and pressure. Multiple trajectory analyses were conducted, including root mean square deviation (RMSD), root mean square fluctuation (RMSF), solvent-accessible surface area (SASA), and radius of gyration (Rg) calculations to assess complex stability, residue-level flexibility, and compactness throughout the simulation period.

### Codon Optimization

Codon optimization of the multi-epitope vaccine construct was performed using the Java Codon Adaptation Tool (JCat) to maximize expression efficiency in Escherichia coli K12 (Grote, Hiller et al. 2005). The Codon Adaptation Index (CAI) and GC content percentage were recorded to ensure optimal expression parameters. The optimized vaccine sequence was cloned into the pET28a (+) expression vector retrieved from the SnapGene official database. Restriction sites XhoI and EcoRI were introduced at the N-terminal and C-terminal regions of the optimized nucleotide sequence to facilitate directional cloning into the vector. The final recombinant plasmid was designed using the SnapGene tool.

### Immune Simulation

The immunogenic performance of the vaccine construct was evaluated using the C-ImmSim server (http://150.146.2.1/C-IMMSIM/), an agent-based computational platform that simulates mammalian immune responses. The server simulates the activity of B cells, T cells, natural killer (NK) cells, macrophages, dendritic cells, and cytokine networks to predict the vaccine’s immunological impact (Rapin, Lund et al. 2011). To reflect diverse human immune backgrounds, HLA alleles A0201, A0101, B0702, B0802, DRB10101, and DRB10301 representing diverse MHC class I and II haplotypes were selected for the simulation. The simulation volume was set to 50 lymphoid compartments, and 1050 time steps were executed.

Immunizations were simulated by antigen injections at time steps 1, 64, and 128 to represent primary immunization and booster doses.

## Results

### Protein Sequence Retrieval

The amino acid sequences encoding Hemagglutinin (HA) and Neuraminidase (NA) from A/swine/Hubei/10/2008 (H10N5) were retrieved from the National Center for Biotechnology Information (NCBI) using the accession numbers AFR42734.1 and AFR42736.1 respectively, and its multiple sequence alignment was performed with homologous proteins of the avian H3N2 strain (HA: JX308801.1, NA: JX286593.1) and human H3N2 strain (HA: OQ350032.1, NA: OQ350034.1) for conservation analysis. Each protein sequence was independently subjected to epitope prediction analyses.

### B-cell Epitopes

Linear B-cell epitopes were predicted separately for both HA and NA proteins of the reference H10N5 strain using computationally validated platforms including ABCpred, BCEpred, and BepiPred, with a threshold of 0.51 and default parameters applied to each protein. Initially, ten epitopes of 16-mer length were predicted for the HA protein, while sixteen 16-mer epitopes were predicted for the NA protein respectively.

The HA-derived epitopes demonstrated antigenicity scores ranging from 0.51 to 1.26, with toxicity prediction support vector machine (SVM) scores between –0.57 and –1.40. In contrast, NA-derived epitopes exhibited antigenicity scores of 0.11 to 0.25 and SVM toxicity scores of 0.41 to 0.85. The complete results of linear B-cell epitope prediction are listed in Table S1.

### CTL Epitopes

Cytotoxic T lymphocyte (CTL) epitopes were predicted independently from HA and NA proteins of H10N5 using NetMHCpan 2.1 with default parameters and a selection threshold of IC50 < 500 nM applied to both proteins. Based on the applied selection criteria, ten CTL epitopes of 15-mer length were identified from the combined HA and NA proteins. HA-derived epitopes exhibited an IC50 value of 95 nM, a percentile rank of 0.04, and a prediction score of 0.85, while NA-derived epitopes demonstrated an IC50 value of 42 nM, a percentile rank of 0.06, and a prediction score of 0.78. The results of CTL epitope prediction for HA and NA genes are presented in Table S2 and S3.

### HTL Epitopes

Helper T lymphocyte (HTL) epitopes, which represent antigenic peptides recognized by CD4+ T cells, were predicted from HA and NA proteins of the H10N5 strain using the NetMHCpan 4.0 server with default parameters. Initially 15 (15 mer) epitopes were predicted for HA while 11 (15 mer) epitopes were predicted for NA protein. Selection criteria included IC50 < 50 nM, prediction score > 0.9, and percentile rank < 0.5. Based on these stringent criteria, five HTL epitopes of 9-mer length were selected from HA and six epitopes of equal length were predicted from the NA protein.

HA-derived HTL epitopes exhibited IC50 values approaching 50 nM, percentile ranks below 0.5, and prediction scores exceeding 0.9. While, the NA-derived HTL epitopes demonstrated comparable binding affinity (IC50 of 42 nM), a percentile rank of 0.06, and a prediction score of 0.78. The predicted epitopes underwent additional validation using VexiJen 3.0, AllerTOP v2.1, and ToxinPred, demonstrating antigenicity scores ranging from 0.51 to 1.26 and toxicity prediction SVM scores ranging from –0.57 to –1.40. The predicted HTL epitopes for HA and NA proteins are listed in Table S4 and S5 respectively.

### Epitope Prediction Criteria

The predicted epitopes were screened based on antigenicity, allergenicity, and toxicity. Epitopes with an antigenicity score ≥ 0.51 in VaxiJen were retained. Conversely, epitopes demonstrating non-allergenicity and non-toxicity in AllerTOP and ToxinPred, respectively, were selected using default parameters.

### Vaccine Construct Design

Epitopes derived from the Hemagglutinin (HA) and Neuraminidase (NA) proteins of the reference strain Influenza A/swine/Hubei/10/2008 (H10N5) were incorporated into the final vaccine construct. To maintain an adequate balance between antigenicity and suitable vaccine size, only top eight B-cell epitopes (seven from HA and one from NA), six MHC-II-restricted HTL epitopes (three from each protein), and six MHC-I-restricted CTL epitopes (one from HA and five from NA). These components were assembled using a strategic linker strategy to prevent epitope fusion and facilitate proper immune processing.

The epitope-to-epitope linkage for both B-cell and HTL epitopes was achieved using the GPGPG linker, while the AAY linker was employed for CTL epitopes. To assemble the final construct, the linker EAAAK was used to sequentially join the selected epitopes, with HTL epitopes linked to CTL epitopes, and subsequently to B-cell epitopes, ensuring structural continuity and stability within the construct. The N-terminal region of the vaccine structure was linked to the adjuvant β-defensin avian through an EAAAK linker, while the C-terminal region was connected to a 6×His tag for purification using the same linker. The linkers serve dual functions of preventing epitope fusion and facilitating proper antigen processing. Following assembly, the finalized construct contained 419 amino acids with a complete one-letter amino acid sequence provided in Figure 1.

**Figure 1:**
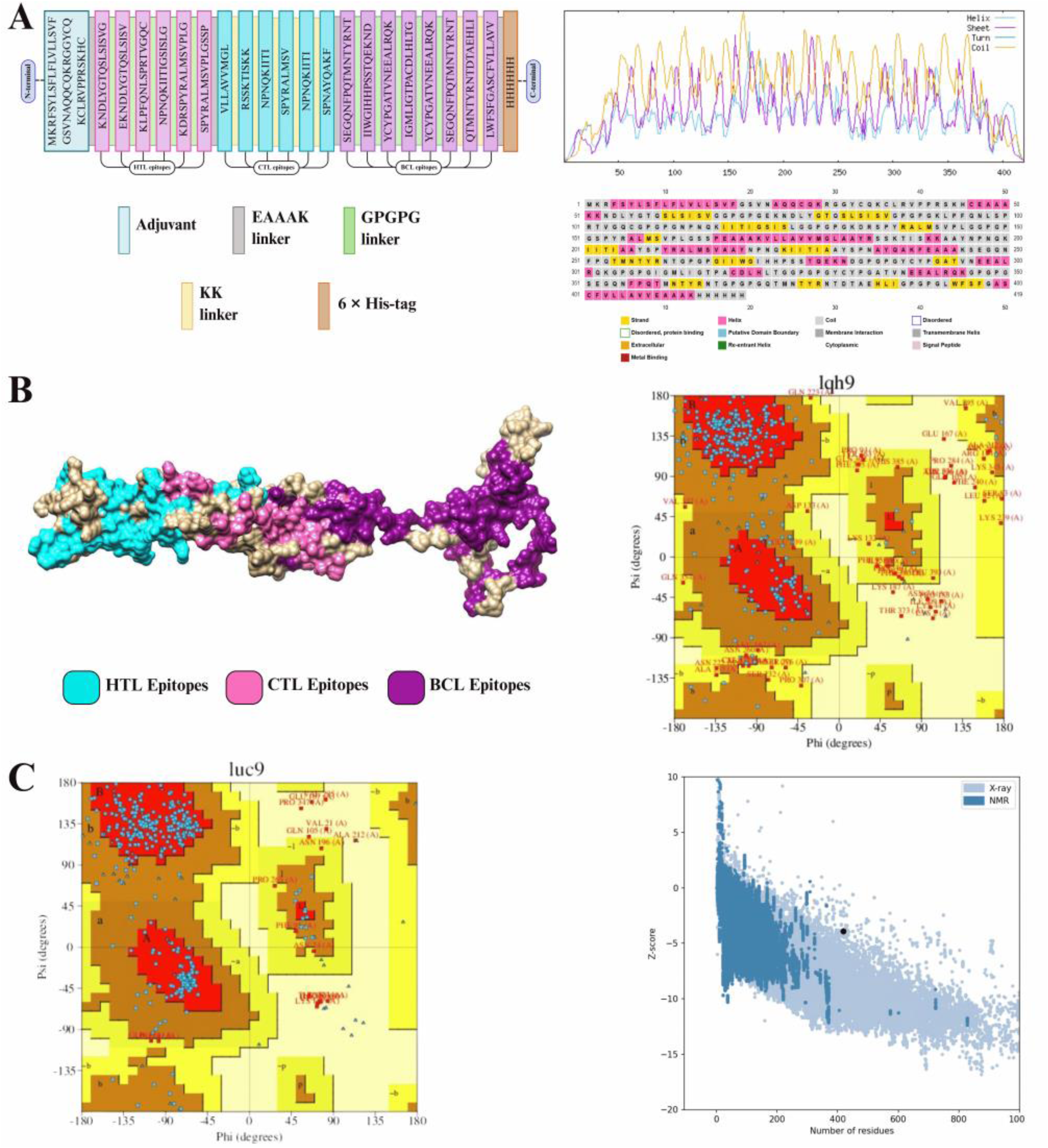
Schematic diagram of the H10N5 strain-based influenza virus vaccine construct and predicted structural features.

### Physicochemical Properties

Physicochemical profile of the vaccine construct was evaluated using Expasy Protparam, to assess its suitability for production and clinical application. The final sequence comprised 419 amino acid residues with a molecular weight of approximately 43.9 kDa, which is compatible with industrial-scale production. The theoretical isoelectric point (pI) was calculated to be 9.50, indicating the construct’s acid-base properties at physiological conditions.

Stability analysis revealed favorable characteristics across multiple systems. The estimated half-life was 30 hours in mammalian reticulocytes (in vitro), greater than 20 hours in yeast (in vitro), and greater than 10 hours in Escherichia coli (in vivo), reflecting good in vivo and in vitro stability across diverse expression systems. The instability index was calculated to be 31.15, further confirming the construct’s stability. The aliphatic index, a measure of thermostability, was determined to be 66.90, while the Grand Average of Hydropathicity (GRAVY) score was 0.363, indicating the construct is slightly hydrophobic. The extinction coefficient was estimated at 41,300 M⁻¹ cm⁻¹.

Analysis of charge distribution revealed important implications for protein behavior. The vaccine construct contained 19 negatively charged residues (aspartate and glutamate) and 36 positively charged residues (arginine and lysine), resulting in a net positive charge at physiological pH. Solubility prediction using SOLPro yielded a probability of 0.763092, indicating favorable aqueous solubility properties suitable for vaccine formulation.

### Secondary Structure Prediction

The secondary structure of the vaccine construct was predicted using SOPMA and PSIPred to understand its functional and structural properties. SOPMA analysis, conducted with a window width of 17, a similarity threshold of 8, and four structural states (helix, sheet, turn, and coil), revealed that the construct consisted of alpha-helices (53.65%), random coils (65.16%), extended beta-strands (19.81%), and beta-turns (2.39%). The high percentage of helical and coil structures suggests a flexible yet structured backbone suitable for immune recognition. PSIPRED analysis further revealed the presence of helices, strand, and coil regions in the vaccine sequence, with results illustrated in Figure 1, confirming the reliability of the SOPMA predictions.

### Tertiary Structure Prediction

The tertiary structure of the vaccine construct was determined through comparative modeling and refinement. Using I-TASSER, five structural models of the vaccine construct were generated and subsequently ranked based on C-score, TM-score, and RMSD values. All five models were then analyzed using the PDBsum server to assess their stereochemical properties and structural quality. Based on Ramachandran plot analysis, the model with 84.4% residues in favored regions and the fewest outliers was selected for further refinement, as this criterion indicates superior stereochemical quality.

The selected model was submitted to GalaxyRefine for optimization through structural adjustment and side-chain repacking, which generated five refined structural models. The best-refined model was identified based on comprehensive quality metrics. This model exhibited a GDT-HA (Global Distance Test-High Accuracy) score of 0.8908, an RMSD (Root Mean Square Deviation) of 0.555 Å, and a Clash score of 12.2. Additionally, the model demonstrated excellent stereochemical quality with only 0.0% poor rotamers and 84.4% of residues positioned within Ramachandran favored regions. The final refined structure was validated using the ProSA-web server, which yielded a Z-score of ∼-4.00, indicating high structural quality. Results of structural refinement are provided in Table S6.

Additional validation via the PROCHECK server confirmed the superior stereochemical properties of the model. Analysis revealed that 58.7% of residues were located in the most favored regions, 18% in additionally allowed regions, 9.8% in generously allowed regions, and only 3.5% in disallowed regions. G-factor analysis further supported the stereochemical reliability of the structure. The overall G-factor was-0.56, while main-chain bond lengths and bond angles yielded G-factors of-1.595 and-2.328, respectively. The dihedral angle distribution (φ/ψ) demonstrated favorable characteristics with a G-factor of-1.49, collectively indicating a reliable and well-modeled three-dimensional structure suitable for functional studies.

### Molecular Docking with TLRs

The vaccine construct was evaluated for its ability to interact with Toll-like receptors (TLRs), which are critical pattern recognition receptors in innate immunity. Specifically, the vaccine was docked with TLR3 and TLR7 receptors. Interaction analysis of the vaccine-TLR3 complex using PDBsum revealed significant interaction features. The complex was stabilized by 2 salt bridges, 16 hydrogen bonds, and 192 non-bonded contacts, with no disulfide bonds contributing to the interface (See Figure 2A). In contrast, the vaccine-TLR7 complex displayed 2 salt bridges, 16 hydrogen bonds, and 213 non-bonded contacts, also without disulfide bond formation (See Figure 2B). The slightly higher number of non-bonded contacts in the vaccine-TLR7 complex suggests that TLR7 forms a more extended interface with the vaccine, indicating higher interaction density compared to TLR3. The results for the PDBSum interaction analysis for the complexes are provided in Table S7 and Table S8 respectively.

**Figure 2:**
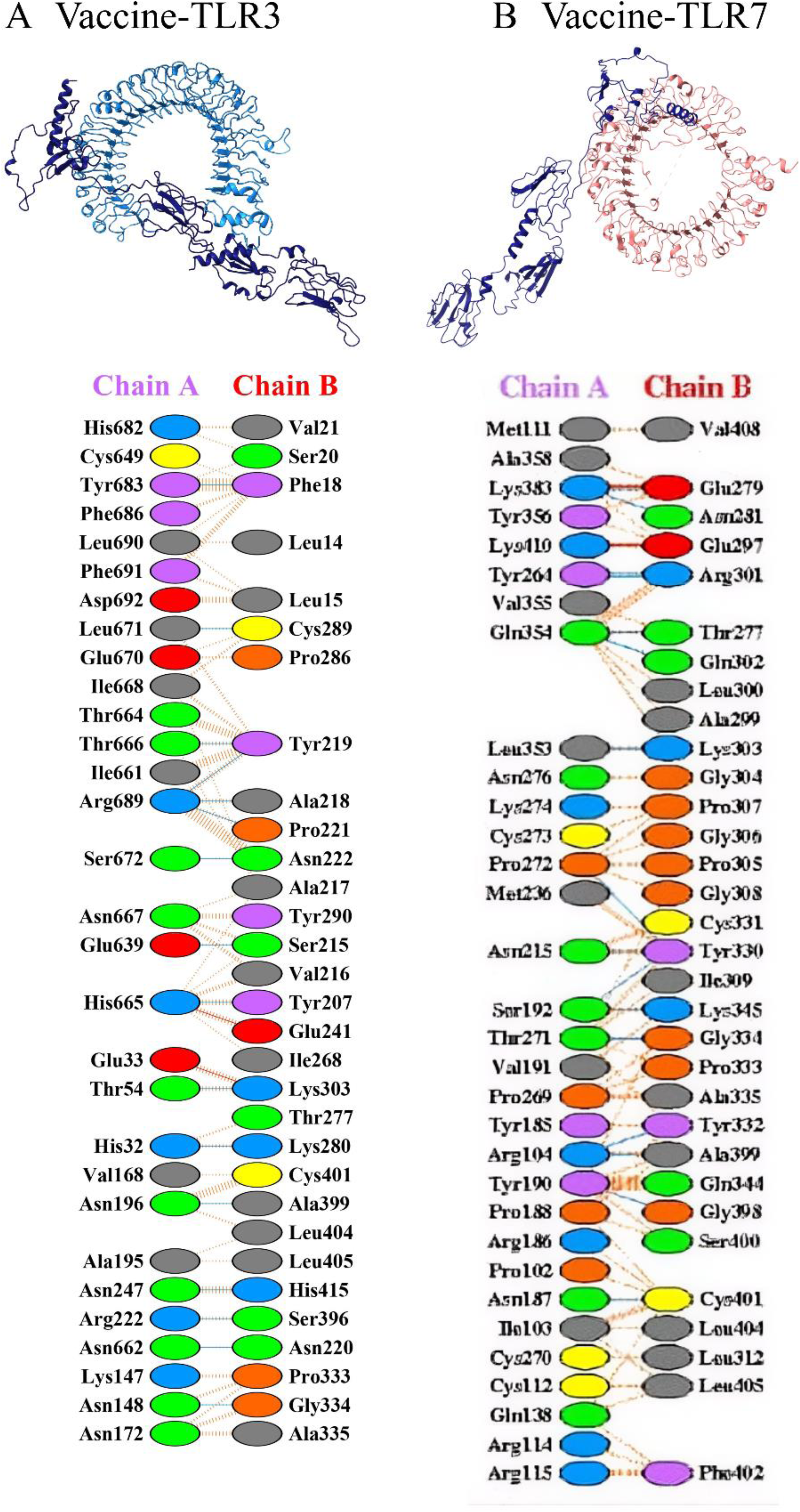
Molecular docking analysis for the influenza H10N5 strain vaccine construct.

Binding affinity analysis performed through the PRODIGY server revealed quantitative differences between the two complexes. The vaccine-TLR3 complex exhibited a binding free energy (ΔG) of –16.1 kcal/mol and a dissociation constant (K_d_) of 1.4 × 10⁻¹² M. In contrast, the vaccine-TLR7 complex showed a ΔG of –16.8 kcal/mol with a dissociation constant (K_d_) of 4.6 × 10⁻¹³ M, indicating that the vaccine-TLR7 complex demonstrated marginally stronger binding affinity based on the lower dissociation constant. Both complexes exhibited highly favorable thermodynamic parameters, suggesting stable and biologically relevant interactions. Results for the PROGIDY analysis are provided in the Table S9 and Table S10 respectively.

Interfacial contact analysis further highlighted subtle but notable differences between the two receptor complexes. In the vaccine-TLR3 complex, the interface was characterized by eight charged-charged interactions, 14 charged-polar interactions, 33 charged-apolar interactions, seven polar-polar interactions, 34 polar-apolar interactions, and 34 apolar-apolar interactions. The non-interacting surface exposed 20.73% charged residues and 39.27% apolar residues. For the vaccine-TLR7 complex, the interface displayed a distinctly different composition with 21 charged-charged interactions, 39 charged-polar interactions, 51 charged-apolar interactions, 12 polar-polar interactions, 22 polar-apolar interactions, and 22 apolar-apolar interactions. The exposed surface at the vaccine-TLR7 interface consisted of 26.75% charged residues and 29.75% apolar residues. These data indicate that the vaccine-TLR7 complex exhibited a relatively higher proportion of charged residue involvement at the interface, whereas the vaccine-TLR3 complex showed greater apolar contributions, reflecting different interaction mechanisms between the vaccine and these two TLR variants.

### Molecular Dynamics Simulation

To evaluate the stability and conformational dynamics of the vaccine receptor complexes during prolonged interaction, molecular dynamics simulations were conducted over extended time scales. The simulations revealed distinct stability trends and dynamic properties between the two complexes.

Root Mean Square Deviation (RMSD) analysis provided insights into overall conformational stability. The vaccine-TLR3 complex stabilized early in the simulation trajectory and maintained RMSD values between 6 and 9 Angstroms (Å) throughout the equilibration and production phases, indicating restricted conformational drift and maintenance of the initial docked configuration (See Figure 3A). In contrast, the vaccine-TLR7 complex displayed substantially higher RMSD values ranging from 22 to 26 Å after equilibration, suggesting more extensive structural rearrangement and accommodation of the vaccine molecule during the dynamic simulations (See Figure 3A). The differential RMSD behavior indicates that while both complexes remained stable, the vaccine-TLR7 interaction involved greater conformational flexibility.

**Figure 3.**
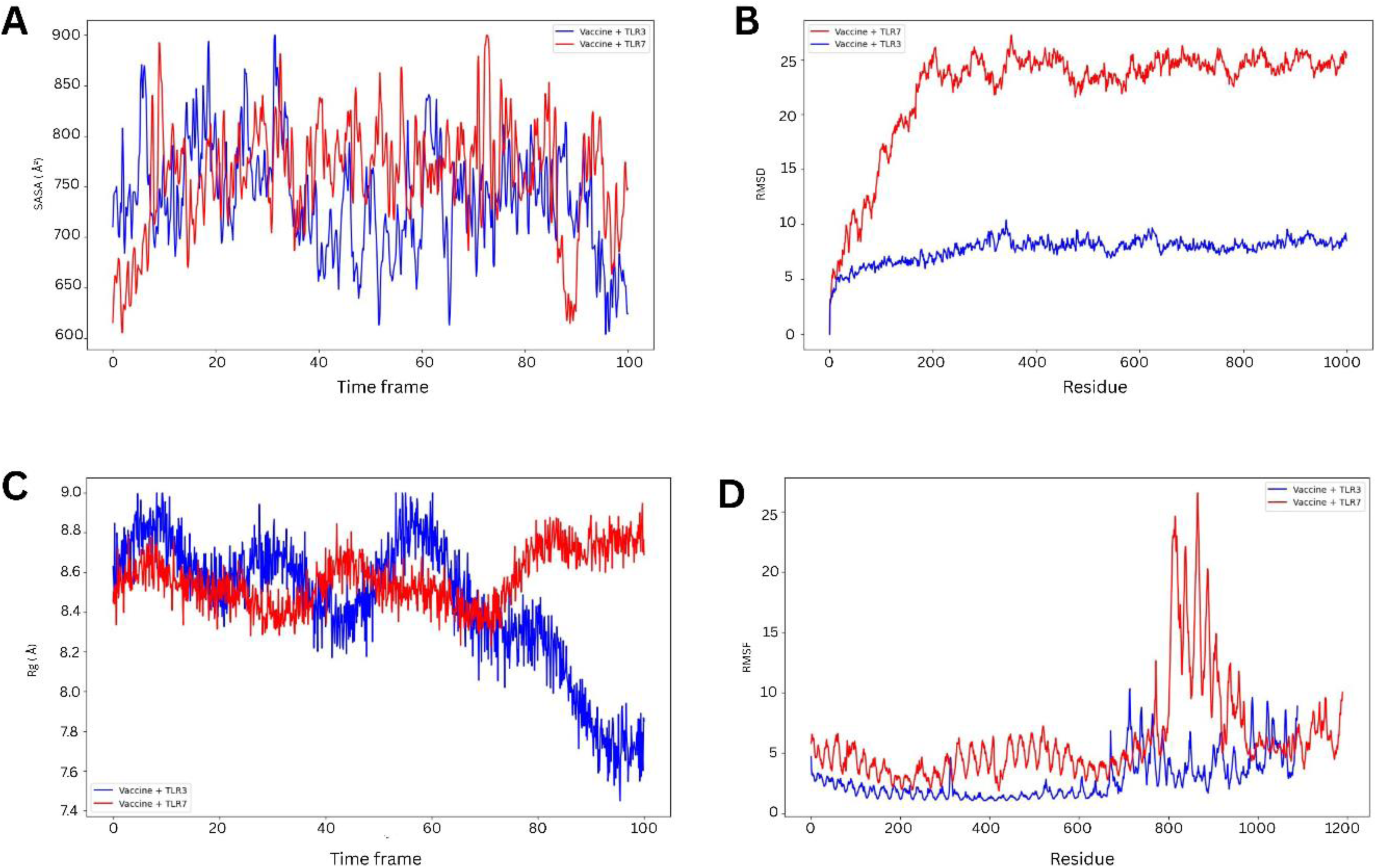
Molecular dynamics simulations for the influenza H10N5 vaccine construct.

Root Mean Square Fluctuation (RMSF) analysis illustrated residue-level fluctuations and revealed localized regions of varying stability. The TLR3 complex exhibited low fluctuations for most residues, with values ranging from 1 to 4 Å, reflecting strong stabilization and reduced mobility upon binding (See Figure 3B). The TLR7-vaccine complex demonstrated pronounced residue-level fluctuations, with a major peak observed between residues approximately 750 to 900, reaching up to 27 Å (See Figure 3B). This elevated fluctuation in the TLR7 complex suggests flexible loop and terminal regions with reduced stabilization compared to the TLR3 complex, potentially indicating regions of the receptor that maintain conformational sampling during the vaccine interaction.

Solvent-Accessible Surface Area (SASA) analysis assessed the hydration status and surface exposure of the complexes throughout the simulation. SASA profiles for both the vaccine-TLR3 and vaccine-TLR7 complexes fluctuated within the 650 to 900 Ų range throughout the simulation trajectory (See Figure 3C). Notably, no major increase in SASA was observed for either complex, indicating the absence of significant structural unfolding or dissociation events, thereby confirming the overall stability of both vaccine receptor interactions under simulated conditions.

Radius of Gyration (Rg) analysis provided information about the spatial compactness of each complex. The TLR3 complex demonstrated progressive compaction, with Rg decreasing gradually from approximately 8.6 Å to approximately 7.7 Å, indicating progressive consolidation and stabilized folding during the simulation (See Figure 3D). The TLR7 complex remained relatively constant between 8.4 and 8.8 Å without appreciable compaction, consistent with the higher RMSD and RMSF values and suggesting lower structural tightness. Collectively, these molecular dynamics data indicate that both complexes maintained structural integrity, but the vaccine-TLR3 interaction achieved greater conformational stability, whereas the vaccine-TLR7 interaction retained greater conformational flexibility.

### Immune Simulation

To evaluate the immunogenicity and immune response profile of the designed vaccine, in silico immune simulation was performed. The vaccine antigen was administered to the simulated host, and the immune response was comprehensively analyzed over the post-immunization period. The results demonstrated a robust and multifaceted immune response characterized by simultaneous activation of humoral and cellular immunity.

The primary antibody response was dominated by immunoglobulin M (IgM), which appeared within 5 to 7 days following antigen administration. The secondary immune response showed significantly elevated levels of IgM, IgG1, and IgM-IgG antibodies, alongside enhanced B-cell proliferation and clonal expansion. The Simpson diversity index analysis indicated broad clonal diversity within the B-cell response, suggesting that the vaccine epitopes effectively engaged multiple B-cell clones, a desirable characteristic for generating comprehensive humoral immunity.

Cellular immune responses were equally robust. Both cytotoxic T cells (Tc) and helper T cells (Th) expanded significantly in the post-immunization period, indicating strong activation of both MHC-I and MHC-II-restricted T-cell responses. Cytokine profiling revealed substantially increased levels of interferon-gamma (IFN-γ) and interleukin-2 (IL-2), confirming the activation of effector T cells and the establishment of a pro-inflammatory response appropriate for combating viral infection.

The innate immune system was also strongly engaged during the vaccine response. Natural killer (NK) cells, which provide critical early antiviral defense, demonstrated rapid expansion following immunization, peaking at 374 cells/mm³ on day 5 post-immunization and subsequently stabilizing at a sustained level of 325 to 345 cells/mm³. Resting dendritic cells (DCs), which serve as critical bridge cells between innate and adaptive immunity, increased from a baseline of approximately 150 cells/mm³ to approximately 200 cells/mm³. Additionally, there was modest activation of antigen-presenting dendritic cell subsets (presenting-1 and presenting-2), indicating the initiation of adaptive immune priming. The coordinated engagement of innate immune effector cells, adaptive B-cell and T-cell responses, and appropriate cytokine milieu collectively demonstrate that the designed vaccine possesses strong immunogenic potential and is likely to generate comprehensive protection against influenza infection.

### Codon Optimization and in silico Cloning

To maximize expression efficiency and optimal protein production in Escherichia coli K12, the designed vaccine protein sequence was codon-optimized using the JCat codon adaptation tool. The optimization process generated a construct comprising nucleotides coding for 419 amino acids with a Codon Adaptation Index (CAI) of 1.0 and a GC content of 55.21%, both values falling within the ideal range for highly efficient E. coli expression. These parameters indicate that the codon usage of the optimized construct matches the host organism’s tRNA availability, ensuring maximal translation efficiency.

Strategic restriction sites were incorporated at both the N-terminal and C-terminal regions of the optimized sequence, ensuring full compatibility with the multiple cloning site of the pET28a (+) vector commonly used for recombinant protein production. Using the SnapGene software tool, the cloning process was performed in silico, generating a recombinant plasmid construct of approximately 6.6 kilobase pairs in total size. This plasmid design provides a complete expression platform for recombinant vaccine production. The results for the in-silico cloning are provided in Figure 4.

**Figure 4.**
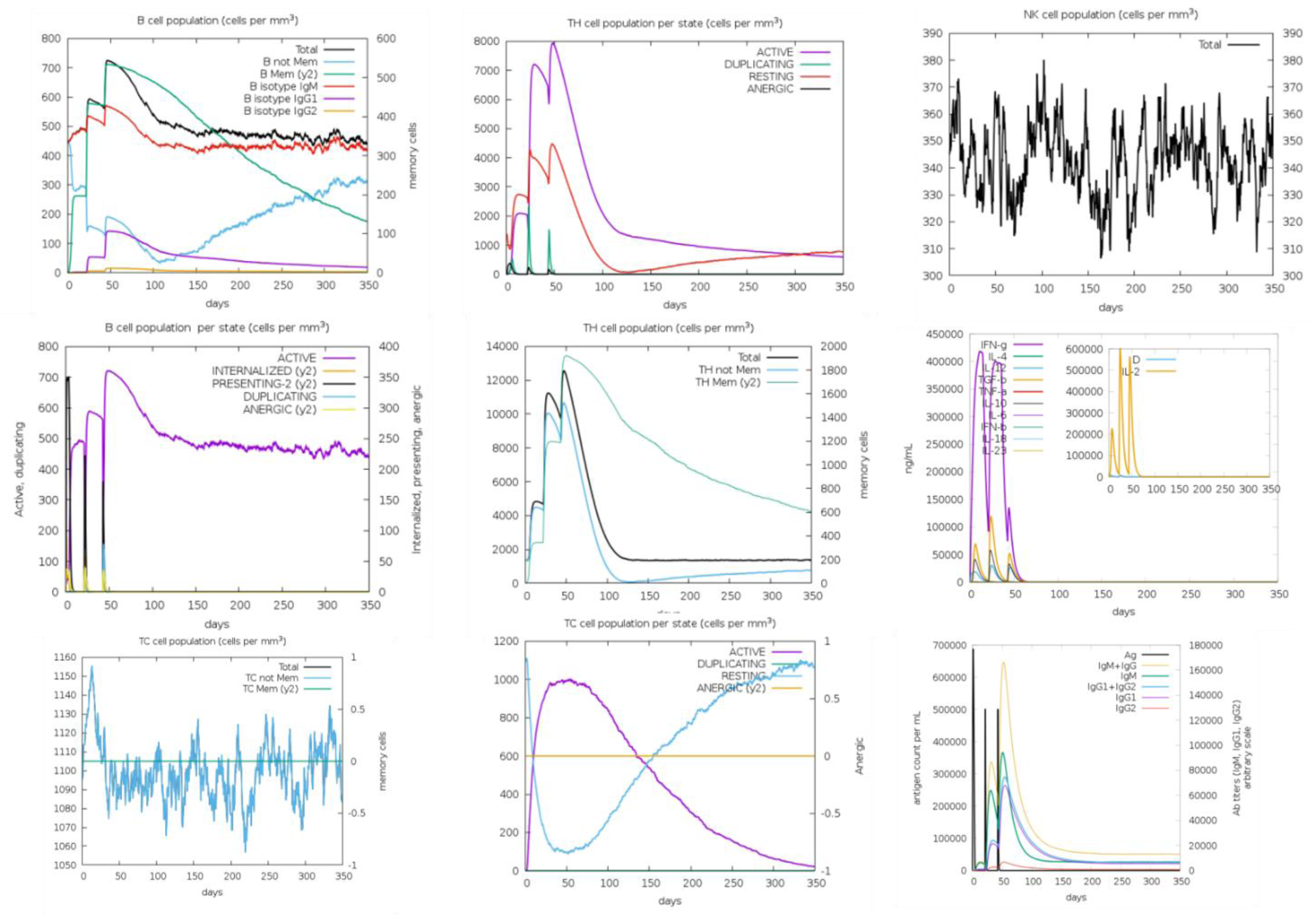
Immune simulations for the influenza H10N5 vaccine construct.

**Figure 5.**
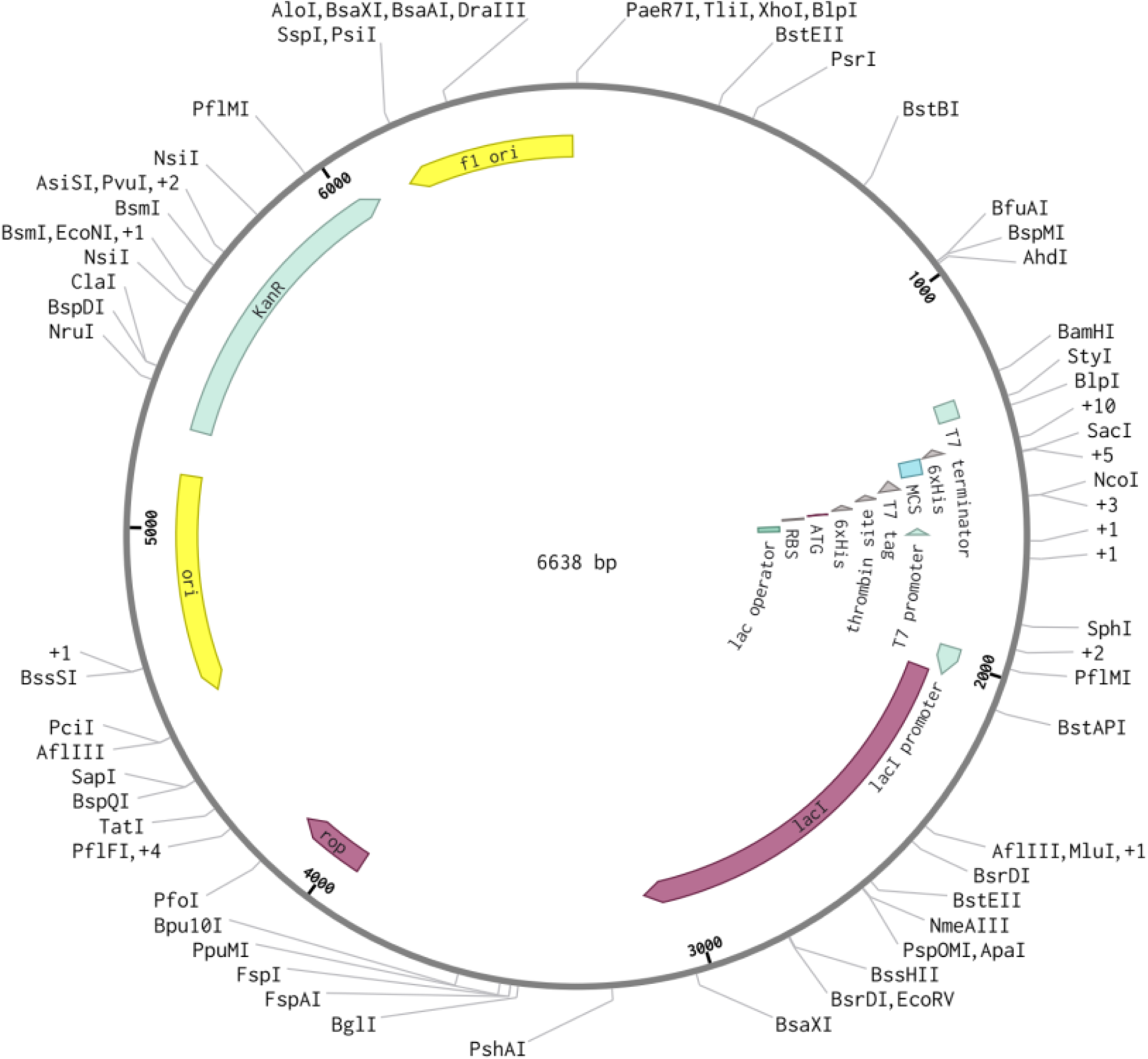
Codon optimization and cloning of Influenza Vaccine Construct.

## Discussion

### Interpreting the Dual H3N2 and H10N5 Targeting Strategy

The fundamental premise of this study was that a single multi-epitope construct could provide immunological coverage against both a dominant seasonal human influenza strain and a potential zoonotic swine subtype that could have mammalian adaptation in the future. The results across structural modeling, docking, molecular dynamics, and immune simulation consistently support this premise, although with qualifications worth examining critically.

The rationale for targeting H3N2 and H10N5 simultaneously was not opportunistic. H3N2 has driven some of the most severe influenza seasons over the past two decades, with disproportionate mortality in elderly individuals arising in part from accumulated antigenic drift that progressively erodes the protective value of pre-existing immunity (Jester, Uyeki et al. 2020). H10N5, in contrast, is an avian-origin virus whose HA gene belongs to the North American lineage, most likely introduced into Asian poultry populations via migratory birds before establishing local circulation through multiple reassortment events incorporating gene segments from H5N2 viruses in domestic waterfowl and H10N7 lineages in wild waterfowl (Yuan, Zhang et al. 2024, Wu, Zhang et al. 2025). The human fatality documented in Anhui Province in January 2024 confirmed first human case infection with one of the H10N5 strain, what phylodynamic data already suggested. this virus has sporadically crossed the species barrier. That it currently lacks known mammalian-adaptation mutations (Yang, Zheng et al. 2025) provides some reassurance but is not grounds for complacency, given the well-established precedent of low-pathogenic avian and swine influenza viruses acquiring pandemic potential through reassortment. Designing against it now, before human-adapted variants emerge, is scientifically prudent.

The multiple sequence alignment between H10N5 and H3N2 HA and NA proteins underpins the entire vaccine strategy. Conservation at the epitope level across these two subtypes is what makes cross-reactive immune priming theoretically achievable. This is not a trivial observation: HA in particular undergoes rapid antigenic drift in seasonal strains, and the identification of genuinely conserved regions shared between an swine/avian low-pathogenic subtype and a human seasonal one required careful alignment across four distinct protein sequences. Epitopes derived exclusively from divergent regions would have generated strain-specific rather than cross-reactive immunity, directly defeating the purpose of a co-protective design.

### Epitope Quality and Multi-Criteria Filtering in Context

The B-cell epitope dataset for HA demonstrated antigenicity scores spanning 0.51 to 1.26 on VaxiJen, a wide range worth examining carefully. Epitopes at the lower threshold near 0.51 represent the minimum bar for predicted antigenicity, and their inclusion in the final construct was justified by confirmed non-toxicity and non-allergenicity rather than antigenicity strength alone. In contrast, NA-derived B-cell epitopes returned antigenicity scores of only 0.11 to 0.25, falling below the 0.51 threshold applied for selection, and were therefore appropriately excluded from the B-cell epitope pool. This is a meaningful finding: NA, while functionally critical as a vaccine target given its role in viral release and its status as an underappreciated correlate of protective immunity (Creytens, Pascha et al. 2021), did not yield strong B-cell epitope candidates under the selection criteria applied here. The construct’s B-cell component therefore draws primarily from HA, while NA contributes more substantially to the CTL and HTL epitope pools. This functional division is biologically defensible but worth stating explicitly, as it shapes the nature of the predicted humoral versus cellular response.

The CTL epitopes from HA exhibited IC50 values of 95 nM with a percentile rank of 0.04, while NA-derived CTL epitopes showed even stronger binding at IC50 42 nM and percentile rank 0.06, both well within the strong-binder threshold of IC50 below 500 nM applied by NetMHCpan 2.1. These values are consistent with strong MHC class I affinity comparable to validated CTL epitopes used in experimental influenza T-cell vaccine studies. The HTL epitope selection was similarly stringent, with IC50 below 50 nM and prediction scores exceeding 0.9 applied as cutoffs, yielding five HA-derived and six NA-derived 9-mer peptides. Taken together, the epitope selection process was multi-layered and conservative, and the final ensemble represents a high-confidence, multi-specificity antigenic portfolio spanning both surface glycoproteins

### Physicochemical Properties

The instability index of 31.15 classifies the construct as stable, a threshold set at values below 40, and this finding is further reinforced by the aliphatic index of 66.90 and estimated half-lives of 30 hours in mammalian reticulocyte systems and over 20 hours in yeast. These are not exceptional values relative to well-characterized recombinant proteins, but they are sufficient to indicate that the construct is unlikely to undergo rapid intracellular degradation before immune processing occurs, which is the functionally relevant criterion.

The GRAVY score of 0.363 warrants careful interpretation. A positive GRAVY score indicates net hydrophobicity, which can be a double-edged property in vaccine design: mild hydrophobicity may facilitate membrane interaction and cellular uptake by antigen-presenting cells, but stronger hydrophobicity creates aggregation risks during recombinant production. At 0.363 the value is mild, and the SOLPro solubility probability of 0.763 provides reassurance that aggregation is unlikely under standard expression conditions. The net positive charge of the construct, with 36 positively charged residues (arginine and lysine) against 19 negatively charged ones, may prove advantageous for electrostatic engagement with negatively charged pattern recognition receptors on the surface of dendritic cells, though this remains speculative in the absence of cell-based binding data.

### Structural Quality: Strengths and Limitations

The final refined model from GalaxyRefine achieved 84.4% Ramachandran-favored residues and a GDT-HA score of 0.8908, both of which are strong indicators of stereochemical quality for a computationally generated multi-domain construct. The RMSD score of 0.555, clash score of 12.2, and poor rotamers of 0.0% fall within the range observed for computationally modeled multi-epitope vaccine structures in the literature, where the artificial junction of heterogeneous peptide sequences with linkers inevitably introduces local geometric irregularities absent from evolved globular proteins (Heo, Park et al. 2013).

Additionally, the PROCHECK analysis identified 3.5% of residues in disallowed Ramachandran regions and a main-chain bond angle G-factor of-2.328, which, while within the range reported in published multi-epitope vaccine models, signals that these regions would benefit from further experimental structural characterization. Atomic-level resolution through X-ray crystallography or cryo-electron microscopy will ultimately be required to confirm whether these computational anomalies correspond to genuine structural tension in the folded protein or are artifacts of the modeling process.

### TLR Docking: Interpreting the Differential Binding Profiles

The binding free energy values obtained through PRODIGY, at-16.1 kcal/mol for TLR3 and - 16.8 kcal/mol for TLR7, sit within the range of strong protein-protein interactions and are comparable to values reported for experimentally validated TLR-ligand complexes. The picomolar dissociation constants (1.4 x 10^-12^ M and 4.6 x 10^-13^ M, respectively) suggest thermodynamically favorable interactions, though it should be noted that PRODIGY-based delta-G estimates are derived from empirical parameterizations of structural interfaces and carry inherent uncertainty, particularly for chimeric proteins not well-represented in the training database. These values are best interpreted as relative rankings between the two complexes rather than absolute thermodynamic measurements.

The vaccine-TLR3 interface was dominated by apolar contacts, with 192 non-bonded contacts and high contributions from charged-apolar (33) and polar-apolar (34) interactions, while the vaccine-TLR7 interface showed a higher proportion of charged-charged (21) and charged-polar (39) interactions among 213 total non-bonded contacts. This distinction maps onto the known structural biology of these two receptors: TLR3 binds double-stranded RNA through a relatively hydrophobic binding cleft on its ectodomain, while TLR7 engages single-stranded RNA through a more electrostatically active recognition pocket (Nafiz, Marlia et al. 2024).

The marginally stronger binding affinity to TLR7 (Kd: 4.6 x 10^-13^ M versus 1.4 x 10^-12^ M for TLR3) is consistent with the known relevance of TLR7 in recognizing single-stranded RNA viruses, of which influenza is a canonical example. This finding suggests the construct may preferentially activate TLR7-mediated MyD88-dependent signaling cascades that drive type I interferon and pro-inflammatory cytokine production, an outcome predicted to enhance the breadth of the subsequent adaptive immune response.

### Molecular Dynamics: Stability Versus Flexibility as Complementary Properties

A key interpretive point from the molecular dynamics data is that the higher RMSD values observed for the vaccine-TLR7 complex, ranging from 22 to 26 Angstroms versus 6 to 9 Angstroms for TLR3, should not be read as evidence of instability. The SASA profiles for both complexes remained stable within 650 to 900 squared Angstroms without significant elevation throughout the simulation. If the vaccine were dissociating from TLR7 or unfolding, SASA would increase markedly. The elevated RMSD and RMSF values, particularly the peak reaching 27 Angstroms at residues 750 to 900, instead reflect conformational sampling in flexible loop and terminal regions of TLR7, a receptor known for structural plasticity in its ectodomain during ligand accommodation.

The progressive Rg compaction in the TLR3 complex, from approximately 8.6 Angstroms to approximately 7.7 Angstroms, is mechanistically distinct, suggesting that the TLR3-vaccine interaction undergoes induced-fit tightening over the simulation timescale consistent with hydrophobic collapse at apolar-dominated protein interfaces. Together, these two complexes exhibit complementary stability modes: conformational lock for TLR3 and dynamic accommodation for TLR7. Both models are biologically viable and have been documented in molecular dynamics studies of TLR-ligand interactions (Sharma, Kumari et al. 2021).

### Immune Simulation

The C-ImmSim simulation results warrant interpretation beyond simple reporting of immune activation. The IgM-dominant primary response transitioning to IgG1 elevation upon booster administration at time steps 64 and 128 follows the expected kinetics of T-dependent B-cell class switching in germinal centers, a response architecture dependent on CD4+ T helper cell activation that the construct’s HTL epitope component is specifically designed to support. The broad Simpson diversity index of the B-cell repertoire is arguably the most clinically significant finding from the immune simulation: it indicates that the vaccine simultaneously engages multiple distinct B-cell clones rather than driving oligoclonal expansion around a single dominant epitope. This property is directly relevant to the problem of influenza immune escape through antigenic drift, as a polyclonal antibody response is far more resistant to single-point mutation-driven evasion than a narrowly focused response against one immunodominant region such as the HA head (Petrova and Russell 2018).

The NK cell peak of 374 cells/mm3 at day 5 post-immunization, stabilizing to 325 to 345 cells/mm3, reflects the early innate antiviral effector component expected from TLR7-driven type I interferon signaling. This temporal pattern is important: NK cell activity in the first week post-exposure provides a critical antiviral bridging defense before antibody and T-cell responses reach protective titers, which typically requires 10 to 14 days. A vaccine that primes this innate effector response is therefore theoretically superior for early-stage infection control compared with constructs that generate only delayed adaptive immunity. Whether this in silico prediction translates to genuine early-phase viral clearance in vivo remains to be tested, but it is a hypothesis directly testable through time-course viral titer measurements in an appropriate animal challenge model.

The robust expansion of both CD8+ cytotoxic T cells and CD4+ helper T cells, accompanied by significantly elevated IFN-gamma and IL-2 production, confirms the activation of a Th1-polarized cellular immune response appropriate for intracellular viral pathogens. IFN-gamma plays a direct antiviral role through upregulation of MHC expression on infected cells, macrophage activation, and direct inhibition of viral replication, while IL-2 drives T-cell clonal expansion and memory formation.

### Adjuvant Selection and Linker Architecture

The incorporation of avian beta-defensin as an N-terminal adjuvant introduces an immunologically active component with well-characterized innate immune stimulatory properties. Beta-defensins function as endogenous host defense peptides that activate dendritic cells, promote Toll-like receptor signaling, and enhance the bridge between innate and adaptive immunity (Herrera-Rodriguez, Meijerhof et al. 2018). The choice of an avian beta-defensin variant is particularly contextually relevant given the avian origin of H10N5, as this adjuvant may be especially effective in potentiating immune responses within the specific context of avian influenza antigen presentation. Its N-terminal placement, connected via the rigid EAAAK linker, ensures physical and functional separation from the epitope array, preventing steric interference with antigen processing while maintaining adjuvant accessibility to innate immune receptors.

The GPGPG linker employed between B-cell and HTL epitopes provides flexible, hinge-like spacing that facilitates independent epitope folding and recognition by B-cell receptors, a design principle supported by multiple computational vaccine studies demonstrating enhanced immunogenicity with flexible linkers (Malonis, Lai et al. 2019). The AAY linker connecting CTL epitopes contributes to efficient proteasomal cleavage and TAP-dependent MHC class I loading, thereby optimizing the presentation of cytotoxic epitopes to CD8+ T cells. This multi-linker architecture reflects a principled approach to vaccine engineering that balances structural integrity with antigen processing efficiency, consistent with established best practices in multi-epitope subunit vaccine design.

### Codon Optimization and Expression Feasibility

The achievement of a Codon Adaptation Index of 1.0 through JCat optimization represents the theoretical maximum for codon harmonization with the Escherichia coli K12 translational machinery, indicating that every codon in the optimized sequence corresponds to the most frequently used codon for each respective amino acid in the host organism. A GC content of 55.21% falls well within the accepted range of 30 to 70% for stable mRNA secondary structure and efficient translation in prokaryotic systems (Grote, Hiller et al. 2005). The successful in silico cloning into the pET28a (+) expression vector provides a complete expression framework translatable to wet laboratory validation, and the incorporation of XhoI and EcoRI restriction sites ensures directional, seamless cloning with minimal risk of reading frame disruption. The resultant 6.6 kb recombinant plasmid size is well within the range manageable by standard bacterial transformation protocols, further supporting the practical feasibility of experimental production.

### Limitations

Several limitations require frank acknowledgment. The H10N5 sequences used originate from a 2008 swine isolate, and the fatal 2024 human case in Anhui Province involved a strain with a distinct reassortment history including segments acquired from avian influenza lineages circulating in Japan, South Korea, Central Asia, and China (Yuan, Zhang et al. 2024). Whether the conserved epitopes identified here are retained in that 2024 lineage must be verified through direct sequence comparison. If key epitopes are absent or mutated in the most current circulating H10N5 strains, the cross-reactive protection predicted by this model may be partial rather than complete. This is not a methodological flaw but rather an inherent limitation of using the only publicly available H10N5 sequences at the time of analysis. The molecular docking employed rigid-body protein-protein methods via ClusPro, which does not model the membrane-proximal endosomal environment where TLR3 and TLR7 actually function; receptor conformation and accessibility in this context may differ from the crystal structures used as input. The C-ImmSim immune simulation, while useful for predicting response direction, cannot capture pre-existing cross-reactive immunity from prior influenza exposure in real-world vaccine recipients, nor individual HLA variability across global populations, two factors known to significantly influence actual vaccine effectiveness (Kunisaki and Janoff 2009). All computational predictions require experimental validation before any translational claims can be substantiated.

### Future Directions

Three concrete experimental priorities follow directly from these findings. First, the recombinant construct should be expressed using the optimized pET28a (+) system described here, with solubility confirmed by size-exclusion chromatography and structural integrity assessed by circular dichroism before immunological testing begins. The CAI of 1.0 and GC content of 55.21% strongly support productive expression, but solubility in practice requires empirical confirmation given the mild hydrophobicity of the construct. Second, TLR3 and TLR7 engagement should be validated using HEK293 reporter cell lines transfected with individual TLR constructs before more complex immunological assays are performed, as this experiment directly tests the most critical mechanistic prediction of the docking analysis at minimal cost.

On the computational side, repeating the multiple sequence alignment and epitope prediction with post-2022 H10N5 sequences, particularly those related to the Anhui 2024 human case, should be a priority before any publication-linked experimental work begins. Population-level HLA coverage analysis using the IEDB Coverage tool across African, Asian, and European cohorts would also strengthen the translational case for this construct as a globally applicable rather than regionally biased vaccine. The inclusion of additional influenza subtypes with documented pandemic potential, particularly H5N1 and H7N9, in the multiple sequence alignment and epitope identification framework could extend the protective breadth of the construct toward a more universal influenza vaccine platform.

As global influenza surveillance continues to detect an expanding diversity of potentially zoonotic subtypes in wild and domestic bird populations across multiple continents, the capacity to rapidly design, computationally validate, and experimentally advance multi-epitope constructs targeting emerging strains will be an increasingly indispensable component of pandemic preparedness (Meseko, Sanicas et al. 2023, Ashraf, Raza et al. 2025). The immunoinformatics pipeline demonstrated here, from sequence retrieval through immune simulation, provides a reproducible and adaptable framework that can be rapidly redeployed against newly emerged influenza variants as sequence data become available. Whether the specific construct reported here advances to experimental and eventually clinical stages will depend on the outcomes of the validation experiments outlined above, but the computational foundation established is rigorous, internally consistent, and designed to be directly actionable.

## Acknowledgments

None.

## Funding

None.

## Conflict of interest

The authors declare no conflicts of interest.

## Ethics approval and consent to participate

This study was entirely in silico, using publicly available data, and did not involve any animal or human participation. Therefore, ethical approval and consent to participate were not required.

## Consent for publication

Not applicable.

## Availability of data

The data supporting the findings of this study are available upon request.

## List of Supplementary Tables

**Supplementary Table S1.**
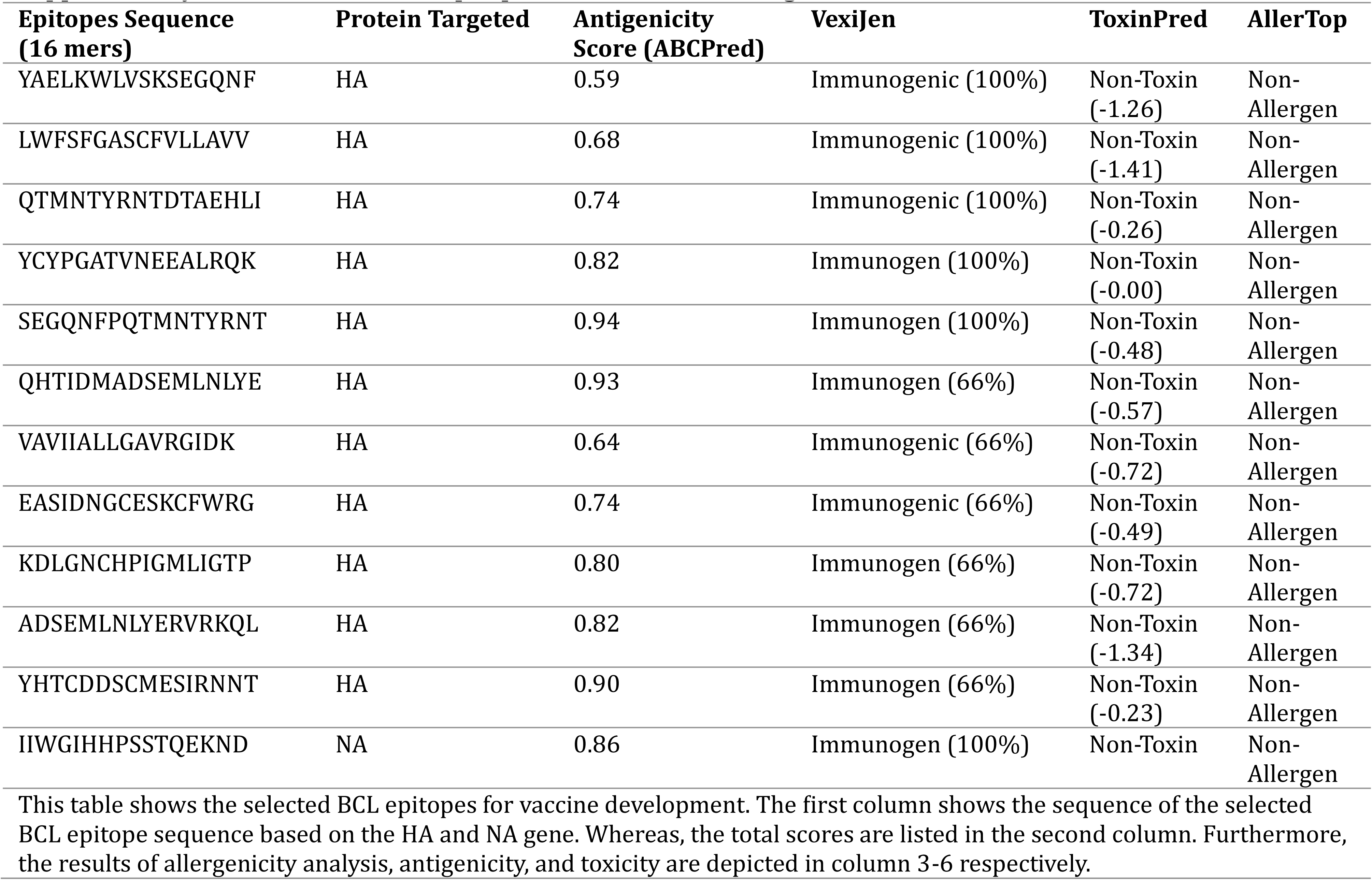
List of BCL epitopes based on HA and NA gene of Influenza Virus strains.

**Supplementary Table S2.**
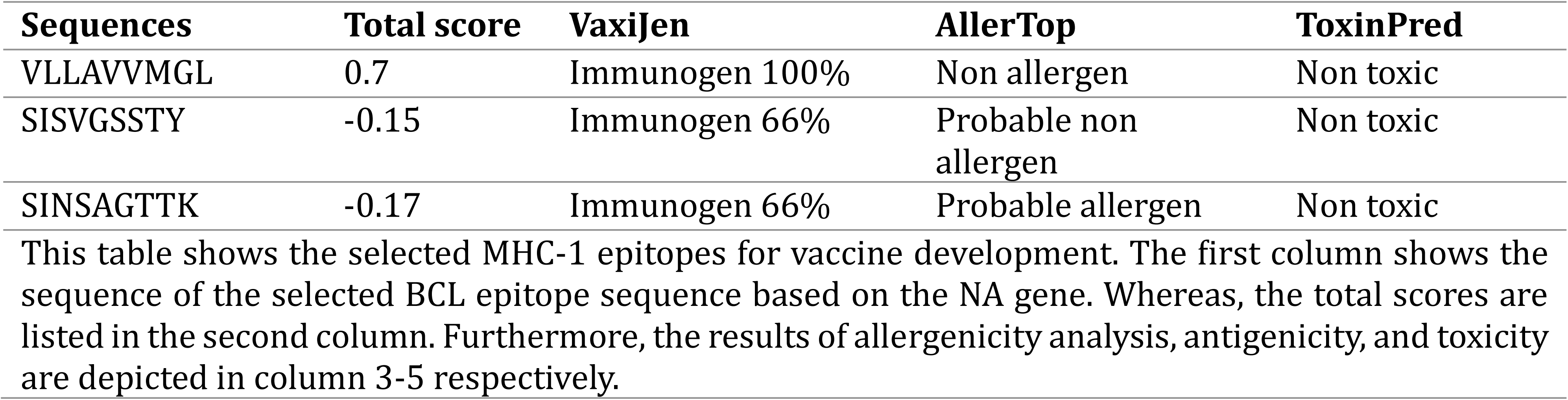
List of CTL epitopes based on the HA gene.

**Supplementary Table S3.**
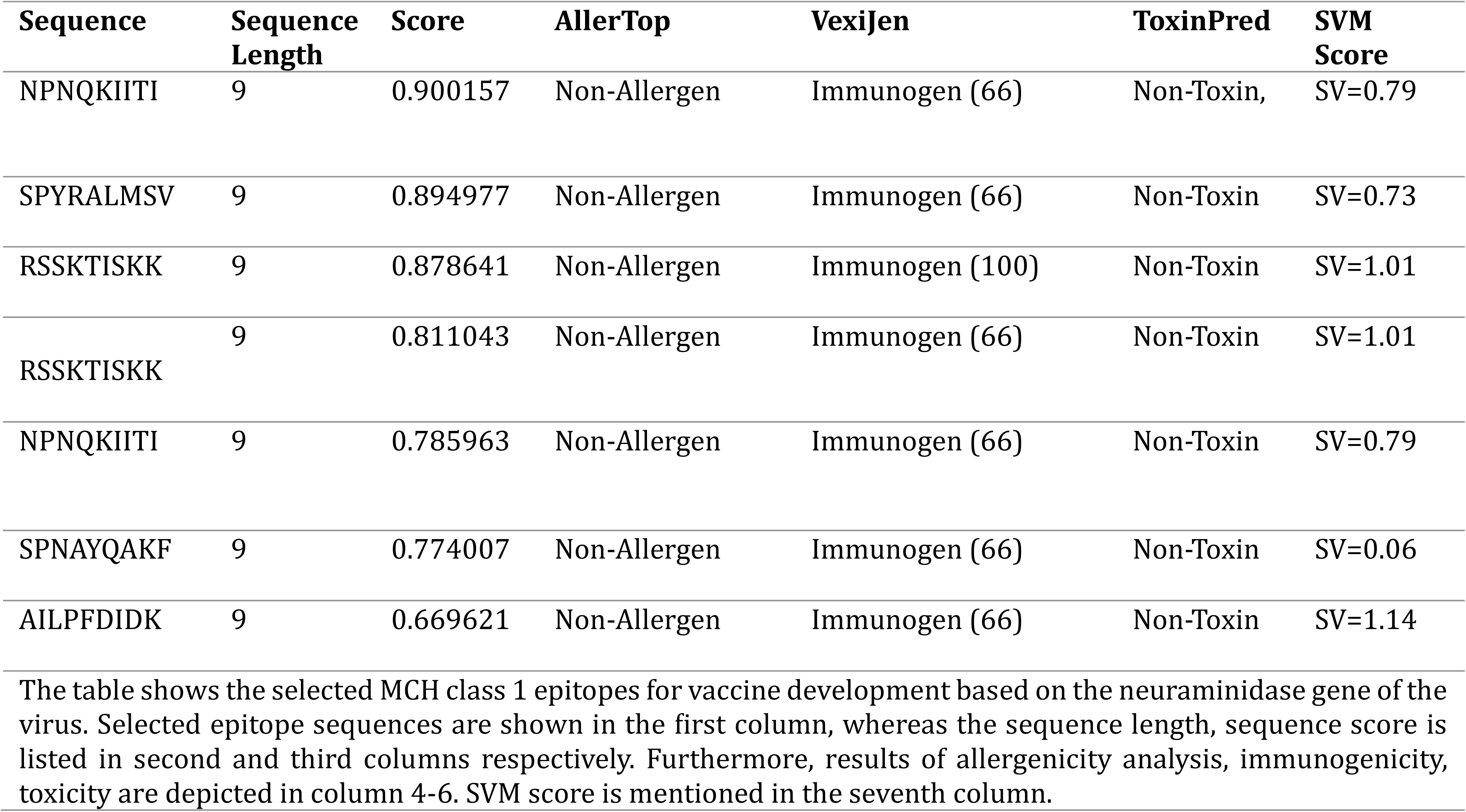
List of CTL Epitopes based on the NA gene.

**Supplementary Table S4.**
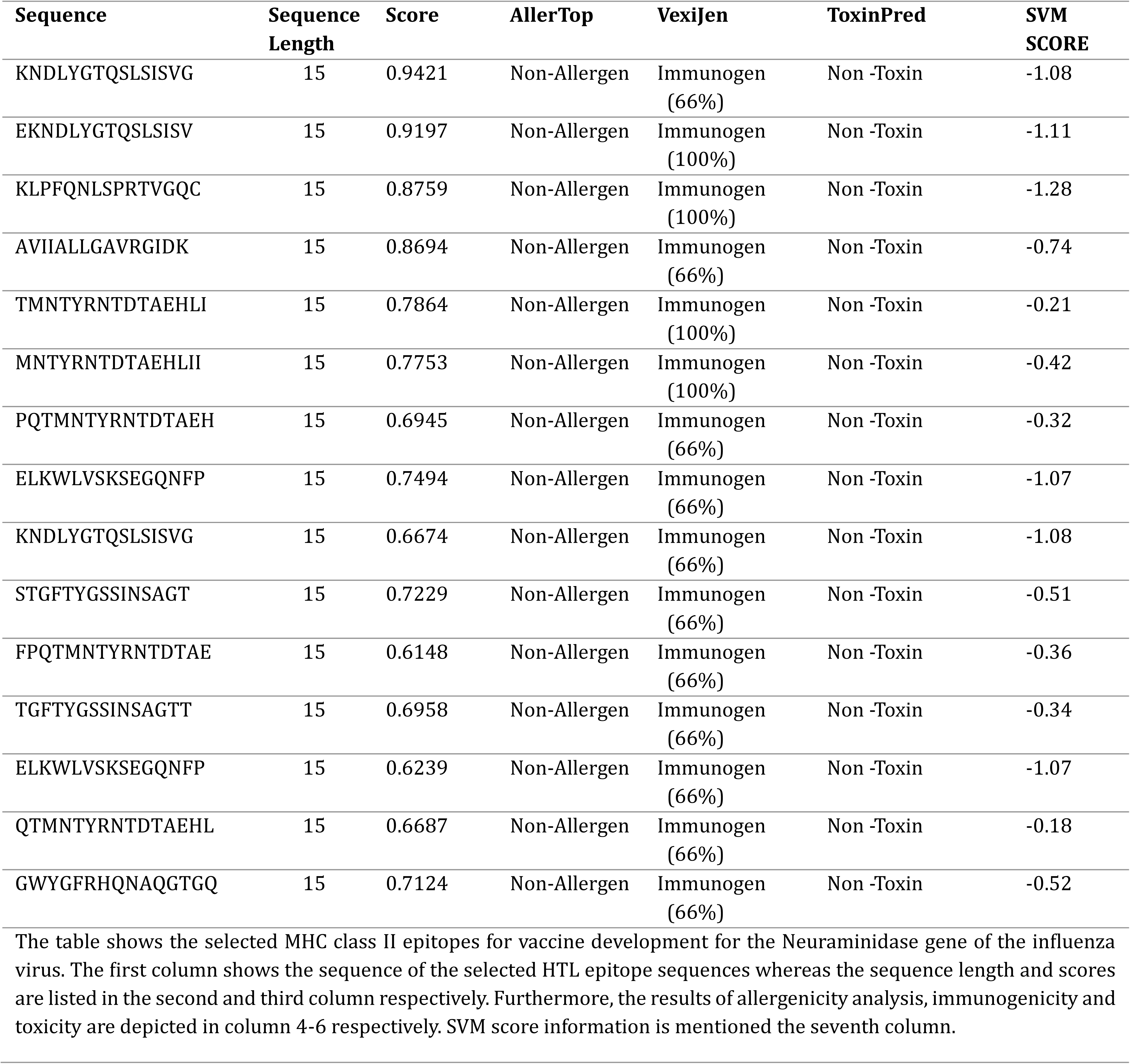
List of selected HTL Epitopes for vaccine construct based on the HA protein.

**Supplementary Table S5.**
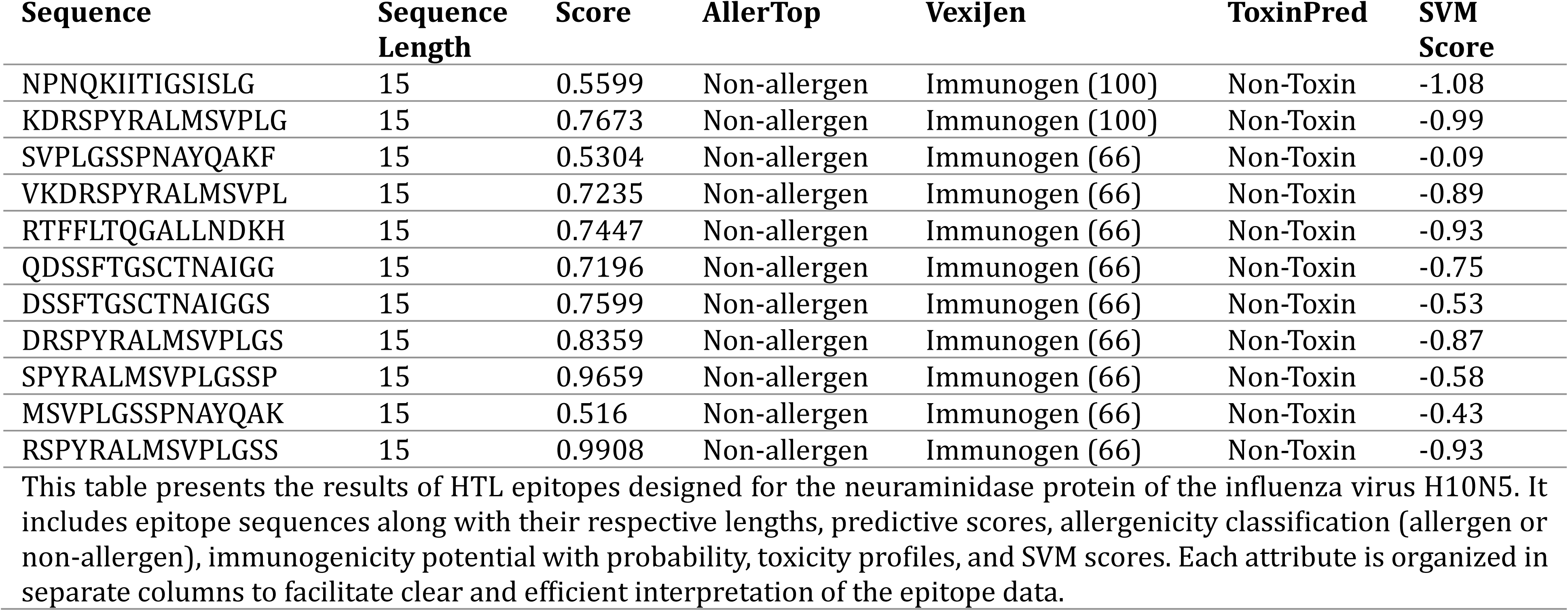
List of Major Histocompatibility Complex (MHC) Class II or HTL Epitopes based on NA gene.

**Supplementary Table S6.**
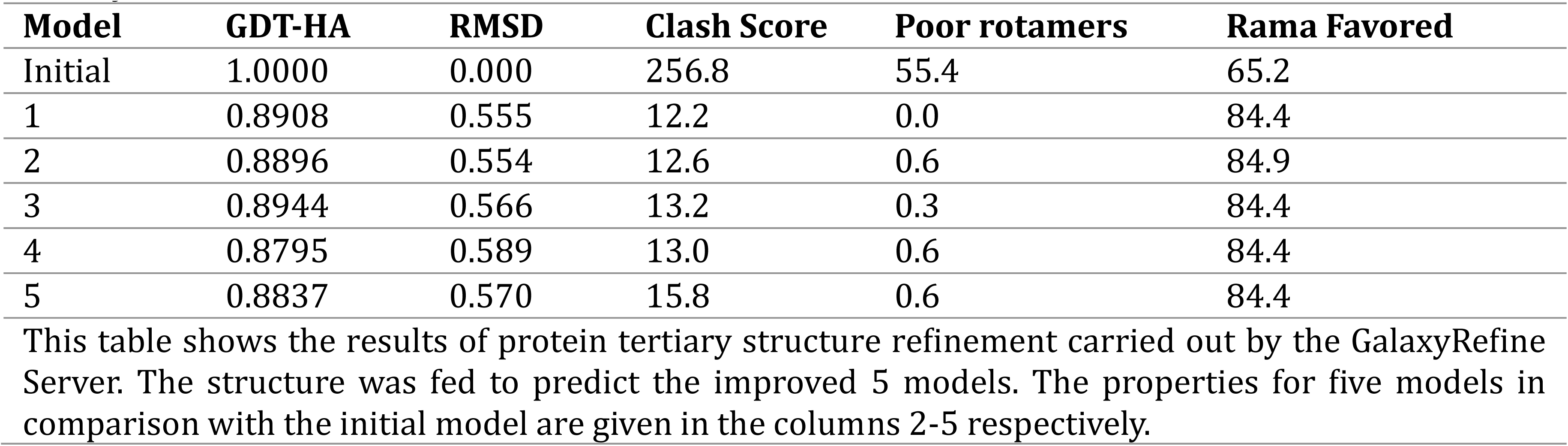
Refinement of Tertiary Structure of the Vaccine Construct Using GalaxyRefine server.

**Supplementary Table S7.**
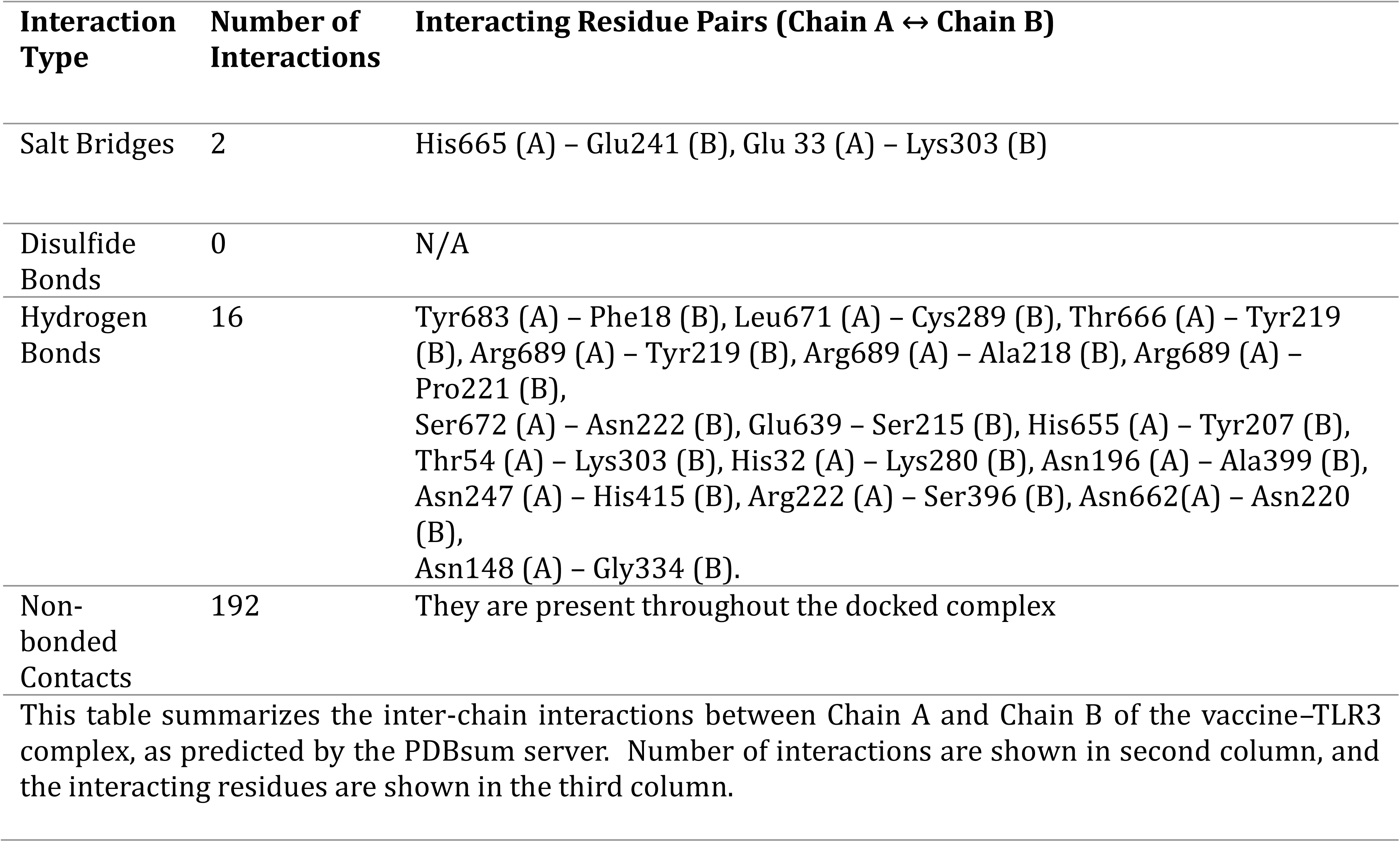
Detailed interaction summary for the vaccine–TLR3 complex.

**Supplementary Table S8.**
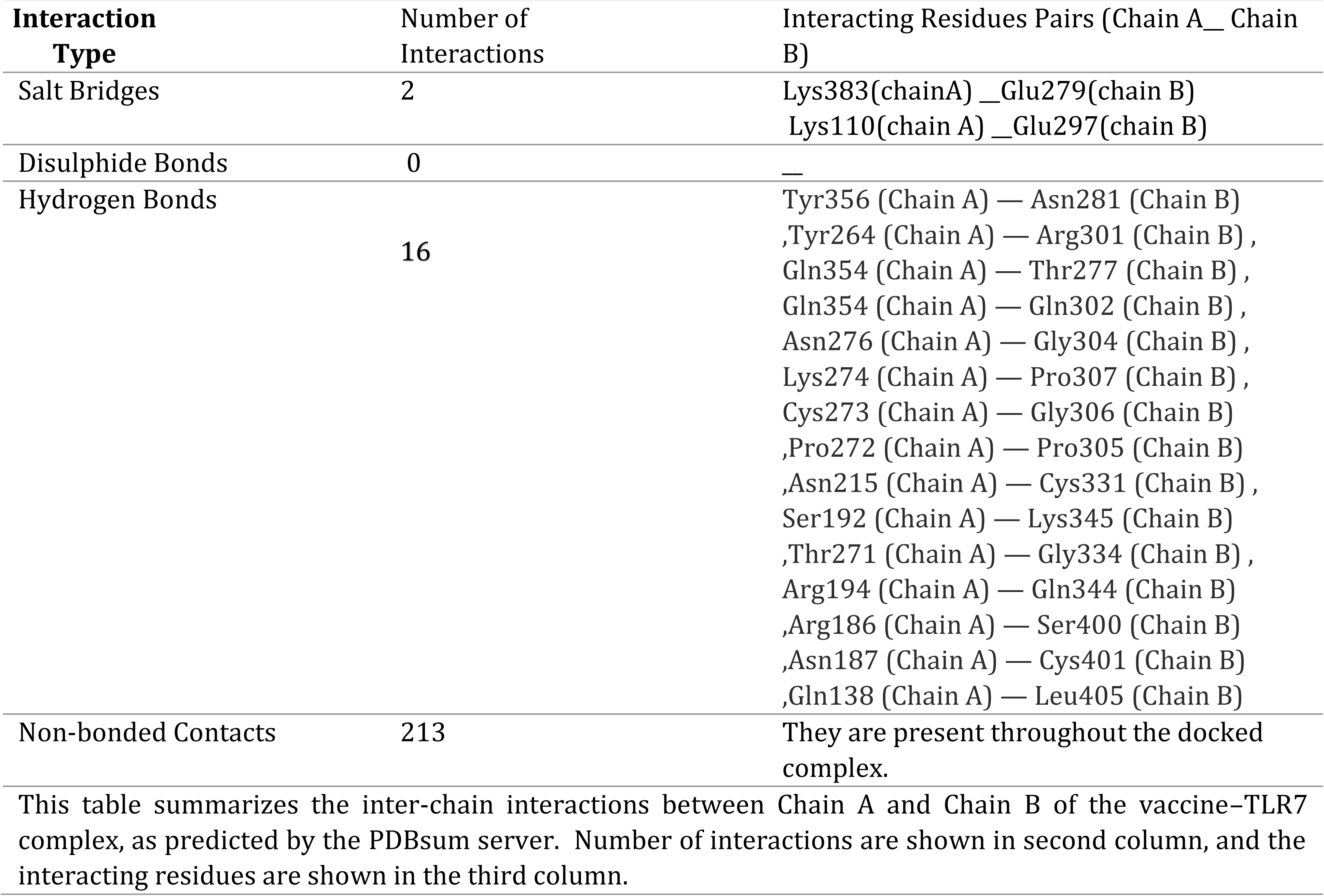
Detailed interaction summary for the vaccine–TLR7 complex.

**Supplementary Table S9.**
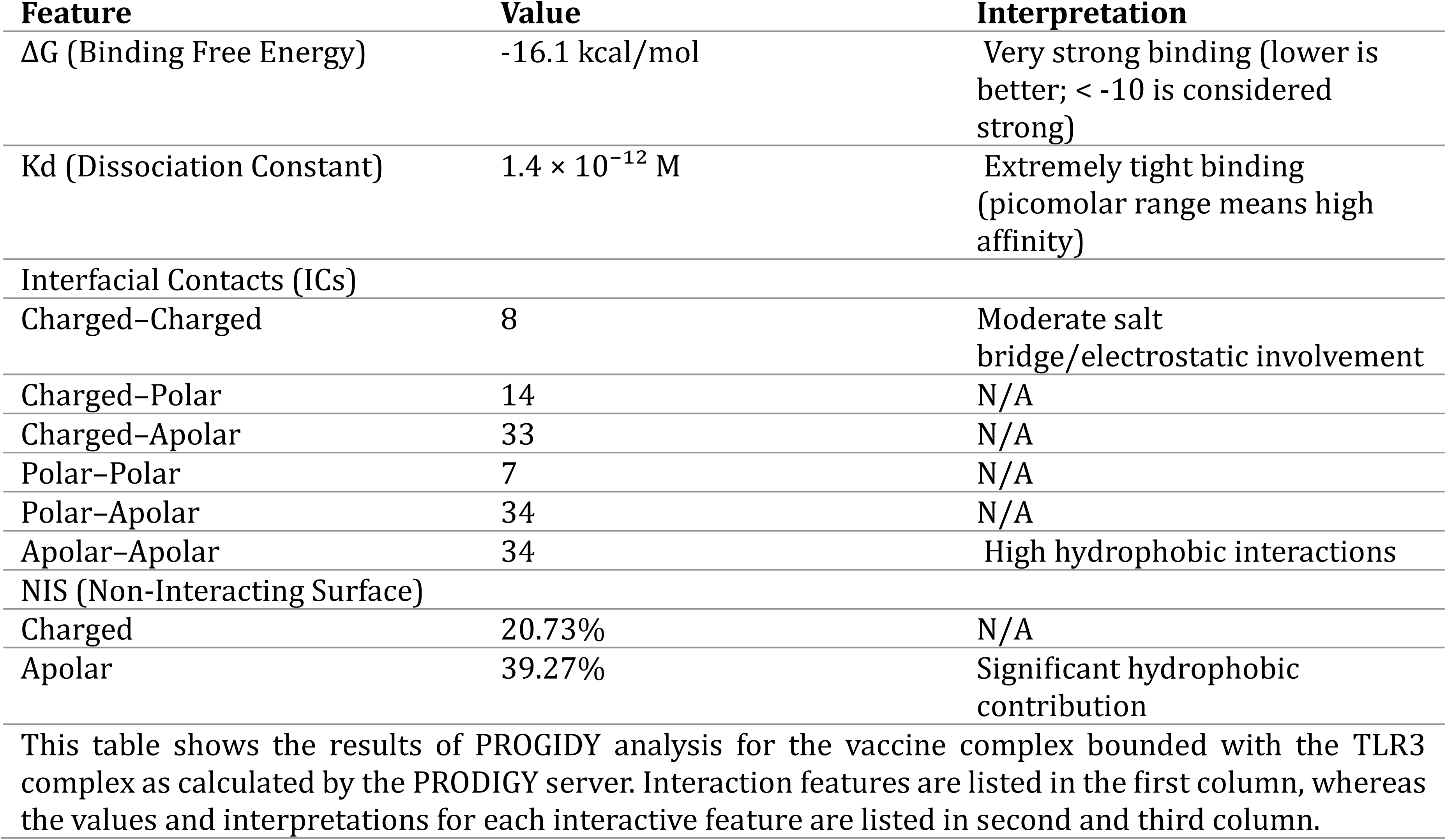
Predicted binding affinity and interaction profile for the vaccine–TLR3 complex.

**Supplementary Table S10.**
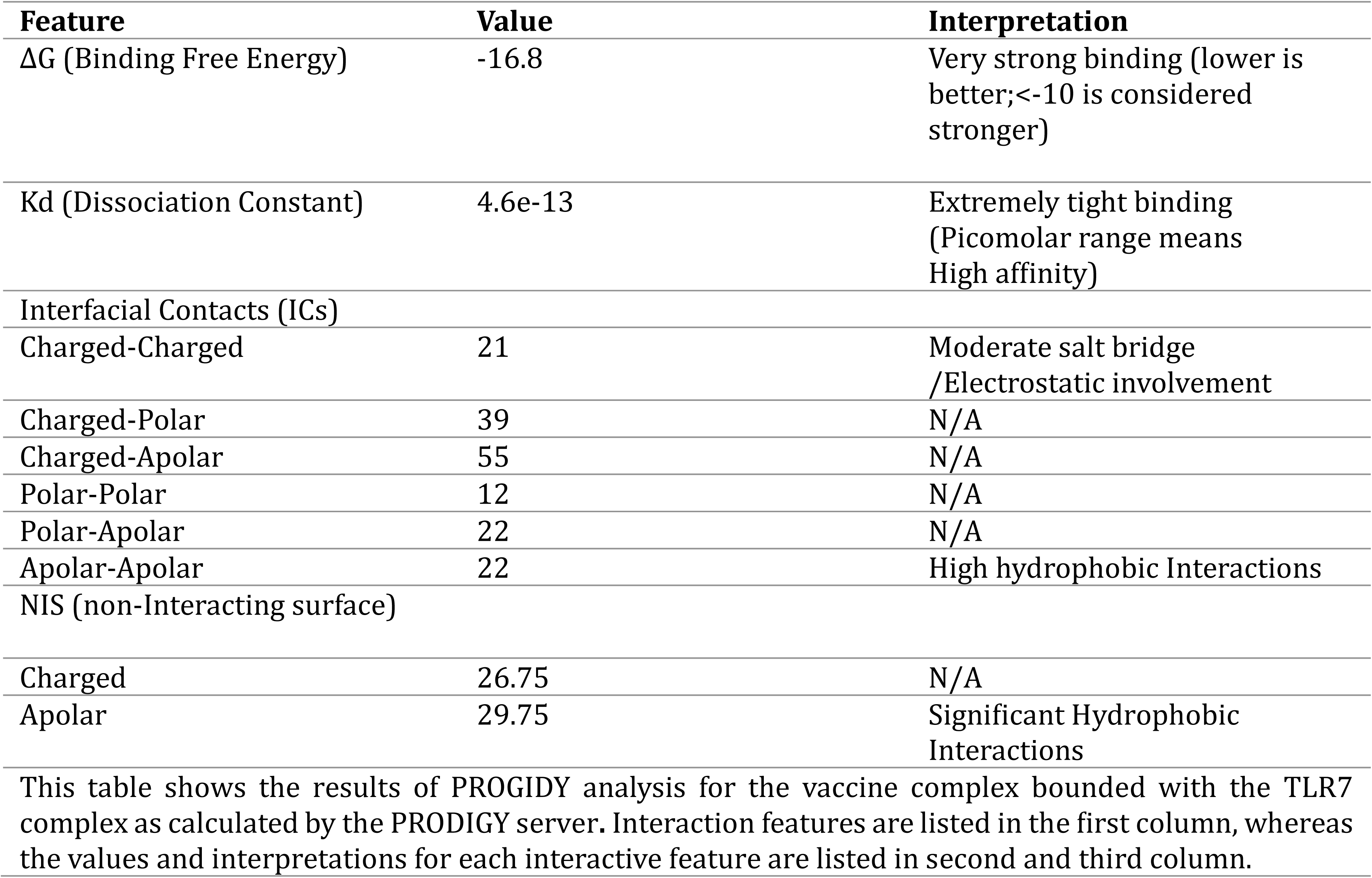
Predicted Binding Affinity and interaction profile for the vaccine and TLR7 complex.

## Notes

### Competing Interest Statement

The authors have declared no competing interest.

