## Supplementary Tables for "Immunoinformatics-Guided Design and In Silico Evaluation of a Multi-Epitope Vaccine Against Influenza A H10N5 and H3N2 Strains Based on Hemagglutinin and Neuraminidase Proteins": Table S1_BCL_Epitopes_MZA_NA_HA.docx

| **Epitopes Sequence**  **(16 mers)** | **Protein Targeted** | **Antigenicity Score (ABCPred)** | **VexiJen** | **ToxinPred** | **AllerTop** |
| --- | --- | --- | --- | --- | --- |
| YAELKWLVSKSEGQNF | HA | 0.59 | Immunogenic (100%) | Non-Toxin  (-1.26) | Non-Allergen |
| LWFSFGASCFVLLAVV | HA | 0.68 | Immunogenic (100%) | Non-Toxin  (-1.41) | Non-Allergen |
| QTMNTYRNTDTAEHLI | HA | 0.74 | Immunogenic (100%) | Non-Toxin  (-0.26) | Non-Allergen |
| YCYPGATVNEEALRQK | HA | 0.82 | Immunogen (100%) | Non-Toxin  (-0.00) | Non-Allergen |
| SEGQNFPQTMNTYRNT | HA | 0.94 | Immunogen (100%) | Non-Toxin  (-0.48) | Non-Allergen |
| QHTIDMADSEMLNLYE | HA | 0.93 | Immunogen (66%) | Non-Toxin  (-0.57) | Non-Allergen |
| VAVIIALLGAVRGIDK | HA | 0.64 | Immunogenic (66%) | Non-Toxin  (-0.72) | Non-Allergen |
| EASIDNGCESKCFWRG | HA | 0.74 | Immunogenic (66%) | Non-Toxin  (-0.49) | Non-Allergen |
| KDLGNCHPIGMLIGTP | HA | 0.80 | Immunogen (66%) | Non-Toxin  (-0.72) | Non-Allergen |
| ADSEMLNLYERVRKQL | HA | 0.82 | Immunogen (66%) | Non-Toxin  (-1.34) | Non-Allergen |
| YHTCDDSCMESIRNNT | HA | 0.90 | Immunogen (66%) | Non-Toxin  (-0.23) | Non-Allergen |
| IIWGIHHPSSTQEKND | NA | 0.86 | Immunogen (100%) | Non-Toxin | Non-Allergen |
| This table shows the selected BCL epitopes for vaccine development. The first column shows the sequence of the selected BCL epitope sequence based on the HA and NA gene. Whereas, the total scores are listed in the second column. Furthermore, the results of allergenicity analysis, antigenicity, and toxicity are depicted in column 3-6 respectively. | | | | | |
