## Supplementary Tables for "Immunoinformatics-Guided Design and In Silico Evaluation of a Multi-Epitope Vaccine Against Influenza A H10N5 and H3N2 Strains Based on Hemagglutinin and Neuraminidase Proteins": Table S2_CTL_HA_MZA.docx

| Table S2. List of CTL epitopes based on the Hemagglutinin gene | | | | |
| --- | --- | --- | --- | --- |
| Sequences | **Total score** | **VaxiJen** | **AllerTop** | **ToxinPred** |
| VLLAVVMGL | 0.7 | Immunogen 100% | Non allergen | Non toxic |
| SISVGSSTY | -0.15 | Immunogen 66% | Probable non allergen | Non toxic |
| SINSAGTTK | -0.17 | Immunogen 66% | Probable allergen | Non toxic |
| This table shows the selected MHC-1 epitopes for vaccine development. The first column shows the sequence of the selected BCL epitope sequence based on the NA gene. Whereas, the total scores are listed in the second column. Furthermore, the results of allergenicity analysis, antigenicity, and toxicity are depicted in column 3-5 respectively. | | | | |
