## Supplementary Tables for "Immunoinformatics-Guided Design and In Silico Evaluation of a Multi-Epitope Vaccine Against Influenza A H10N5 and H3N2 Strains Based on Hemagglutinin and Neuraminidase Proteins": Table S3_CTL Epitopes_MZA_Neuraminidase Gene.docx

| Table S3. List of CTL Epitopes based on the Neuraminidase gene | | | | | | |
| --- | --- | --- | --- | --- | --- | --- |
| Sequence | Sequence  Length | Score | AllerTop | VexiJen | ToxinPred | SVM Score |
| NPNQKIITI | 9 | 0.900157 | Non-ALLERGEN | IMMUNOGEN (66) | Non-Toxin, | SV=0.79 |
| SPYRALMSV | 9 | 0.894977 | Non- ALLERGEN | IMMUNOGEN (66) | Non-Toxin | SV=0.73 |
| RSSKTISKK | 9 | 0.878641 | Non-ALLERGEN | IMMUNOGEN (100) | Non-Toxin | SV=1.01 |
| RSSKTISKK | 9 | 0.811043 | Non-ALLERGEN | IMMUNOGEN (66) | Non-Toxin | SV=1.01 |
| NPNQKIITI | 9 | 0.785963 | Non-ALLERGEN | IMMUNOGEN (66) | Non-Toxin | SV=0.79 |
| SPNAYQAKF | 9 | 0.774007 | Non-ALLERGEN | IMMUNOGEN (66) | Non-Toxin | SV=0.06 |
| AILPFDIDK | 9 | 0.669621 | Non-ALLERGEN | IMMUNOGEN (66) | Non-Toxin | SV=1.14 |
| The table shows the selected MCH class 1 epitopes for vaccine development based on the neuraminidase gene of the virus. Selected epitope sequences are shown in the first column, whereas the sequence length, sequence score is listed in second and third columns respectively. Furthermore, results of allergenicity analysis, immunogenicity, toxicity are depicted in column 4-6. SVM score is mentioned in the seventh column. | | | | | | |
