## Supplementary Tables for "Immunoinformatics-Guided Design and In Silico Evaluation of a Multi-Epitope Vaccine Against Influenza A H10N5 and H3N2 Strains Based on Hemagglutinin and Neuraminidase Proteins": Table S4_HTL Epitopes_HA_MZA.docx

| Table S4. List of selected HTL Epitopes for vaccine construct based on the HA protein | | | | | | |
| --- | --- | --- | --- | --- | --- | --- |
| Sequence | **Sequence**  **Length** | **Score** | **AllerTop** | **VexiJen** | **ToxinPred** | **SVM**  **SCORE** |
| KNDLYGTQSLSISVG | 15 | 0.9421 | Non-Allergen | Immunogen  (66%) | Non -Toxin | -1.08 |
| EKNDLYGTQSLSISV | 15 | 0.9197 | Non-Allergen | Immunogen  (100%) | Non -Toxin | -1.11 |
| KLPFQNLSPRTVGQC | 15 | 0.8759 | Non-Allergen | Immunogen  (100%) | Non -Toxin | -1.28 |
| AVIIALLGAVRGIDK | 15 | 0.8694 | Non-Allergen | Immunogen  (66%) | Non -Toxin | -0.74 |
| TMNTYRNTDTAEHLI | 15 | 0.7864 | Non-Allergen | Immunogen  (100%) | Non -Toxin | -0.21 |
| MNTYRNTDTAEHLII | 15 | 0.7753 | Non-Allergen | Immunogen  (100%) | Non -Toxin | -0.42 |
| PQTMNTYRNTDTAEH | 15 | 0.6945 | Non-Allergen | Immunogen  (66%) | Non -Toxin | -0.32 |
| ELKWLVSKSEGQNFP | 15 | 0.7494 | Non-Allergen | Immunogen  (66%) | Non -Toxin | -1.07 |
| KNDLYGTQSLSISVG | 15 | 0.6674 | Non-Allergen | Immunogen  (66%) | Non -Toxin | -1.08 |
| STGFTYGSSINSAGT | 15 | 0.7229 | Non-Allergen | Immunogen  (66%) | Non -Toxin | -0.51 |
| FPQTMNTYRNTDTAE | 15 | 0.6148 | Non-Allergen | Immunogen  (66%) | Non -Toxin | -0.36 |
| TGFTYGSSINSAGTT | 15 | 0.6958 | Non-Allergen | Immunogen  (66%) | Non -Toxin | -0.34 |
| ELKWLVSKSEGQNFP | 15 | 0.6239 | Non-Allergen | Immunogen  (66%) | Non -Toxin | -1.07 |
| QTMNTYRNTDTAEHL | 15 | 0.6687 | Non-Allergen | Immunogen  (66%) | Non -Toxin | -0.18 |
| GWYGFRHQNAQGTGQ | 15 | 0.7124 | Non-Allergen | Immunogen  (66%) | Non -Toxin | -0.52 |
| The table shows the selected MHC class II epitopes for vaccine development for the Neuraminidase gene of the influenza virus. The first column shows the sequence of the selected HTL epitope sequences whereas the sequence length and scores are listed in the second and third column respectively. Furthermore, the results of allergenicity analysis, immunogenicity and toxicity are depicted in column 4-6 respectively. SVM score information is mentioned the seventh column. | | | | | | |
