## Supplementary Tables for "Immunoinformatics-Guided Design and In Silico Evaluation of a Multi-Epitope Vaccine Against Influenza A H10N5 and H3N2 Strains Based on Hemagglutinin and Neuraminidase Proteins": Table S5_HTL_Epitopes_NA_MZA.docx

| Table S6. List of Major Histocompatibility Complex (MHC) Class II or HTL Epitopes based on NA gene | | | | | | |
| --- | --- | --- | --- | --- | --- | --- |
| Sequence | **Sequence**  **Length** | **Score** | **AllerTop** | **VexiJen** | **ToxinPred** | **SVM Score** |
| NPNQKIITIGSISLG | 15 | 0.5599 | Non-allergen | Immunogen (100) | Non-Toxin | -1.08 |
| KDRSPYRALMSVPLG | 15 | 0.7673 | Non-allergen | Immunogen (100) | Non-Toxin | -0.99 |
| SVPLGSSPNAYQAKF | 15 | 0.5304 | Non-allergen | Immunogen (66) | Non-Toxin | -0.09 |
| VKDRSPYRALMSVPL | 15 | 0.7235 | Non-allergen | Immunogen (66) | Non-Toxin | -0.89 |
| RTFFLTQGALLNDKH | 15 | 0.7447 | Non-allergen | Immunogen (66) | Non-Toxin | -0.93 |
| QDSSFTGSCTNAIGG | 15 | 0.7196 | Non-allergen | Immunogen (66) | Non-Toxin | -0.75 |
| DSSFTGSCTNAIGGS | 15 | 0.7599 | Non-allergen | Immunogen (66) | Non-Toxin | -0.53 |
| DRSPYRALMSVPLGS | 15 | 0.8359 | Non-allergen | Immunogen (66) | Non-Toxin | -0.87 |
| SPYRALMSVPLGSSP | 15 | 0.9659 | Non-allergen | Immunogen (66) | Non-Toxin | -0.58 |
| MSVPLGSSPNAYQAK | 15 | 0.516 | Non-allergen | Immunogen (66) | Non-Toxin | -0.43 |
| RSPYRALMSVPLGSS | 15 | 0.9908 | Non-allergen | Immunogen (66) | Non-Toxin | -0.93 |
| This table presents the results of HTL epitopes designed for the neuraminidase protein of the influenza virus H10N5. It includes epitope sequences along with their respective lengths, predictive scores, allergenicity classification (allergen or non-allergen), immunogenicity potential with probability, toxicity profiles, and SVM scores. Each attribute is organized in separate columns to facilitate clear and efficient interpretation of the epitope data. | | | | | | |
