## Supplementary Tables for "Immunoinformatics-Guided Design and In Silico Evaluation of a Multi-Epitope Vaccine Against Influenza A H10N5 and H3N2 Strains Based on Hemagglutinin and Neuraminidase Proteins": Table S6_MZA.docx

| Table S6. Refinement of Tertiary Structure of the Vaccine Construct Using GalaxyRefine server | | | | | |
| --- | --- | --- | --- | --- | --- |
| Model | **GDT-HA** | **RMSD** | **Clash Score** | **Poor rotamers** | **Rama Favored** |
| Initial | 1.0000 | 0.000 | 256.8 | 55.4 | 65.2 |
| 1 | 0.8908 | 0.555 | 12.2 | 0.0 | 84.4 |
| 2 | 0.8896 | 0.554 | 12.6 | 0.6 | 84.9 |
| 3 | 0.8944 | 0.566 | 13.2 | 0.3 | 84.4 |
| 4 | 0.8795 | 0.589 | 13.0 | 0.6 | 84.4 |
| 5 | 0.8837 | 0.570 | 15.8 | 0.6 | 84.4 |
| This table shows the results of protein tertiary structure refinement carried out by the GalaxyRefine Server. The structure was fed to predict the improved 5 models. The properties for five models in comparison with the initial model are given in the columns 2-5 respectively. | | | | | |
