## Supplementary Tables for "Immunoinformatics-Guided Design and In Silico Evaluation of a Multi-Epitope Vaccine Against Influenza A H10N5 and H3N2 Strains Based on Hemagglutinin and Neuraminidase Proteins": Table S7_PBDSUM_TLR3_VACCINE.docx

| Table S7. Detailed interaction summary for the vaccine–TLR3 complex | | |
| --- | --- | --- |
| Interaction Type | **Number of Interactions** | **Interacting Residue Pairs (Chain A ↔ Chain B)** |
| Salt Bridges | 2 | His665 (A) – Glu241 (B), Glu 33 (A) – Lys303 (B) |
| Disulfide Bonds | 0 | N/A |
| Hydrogen Bonds | 16 | Tyr683 (A) – Phe18 (B), Leu671 (A) – Cys289 (B), Thr666 (A) – Tyr219 (B), Arg689 (A) – Tyr219 (B), Arg689 (A) – Ala218 (B), Arg689 (A) – Pro221 (B),  Ser672 (A) – Asn222 (B), Glu639 – Ser215 (B), His655 (A) – Tyr207 (B),  Thr54 (A) – Lys303 (B), His32 (A) – Lys280 (B), Asn196 (A) – Ala399 (B),  Asn247 (A) – His415 (B), Arg222 (A) – Ser396 (B), Asn662(A) – Asn220 (B),  Asn148 (A) – Gly334 (B). |
| Non-bonded Contacts | 192 | They are present throughout the docked complex |
| This table summarizes the inter-chain interactions between Chain A and Chain B of the vaccine–TLR3 complex, as predicted by the PDBsum server. Number of interactions are shown in second column, and the interacting residues are shown in the third column. | | |
