## Supplementary Tables for "Immunoinformatics-Guided Design and In Silico Evaluation of a Multi-Epitope Vaccine Against Influenza A H10N5 and H3N2 Strains Based on Hemagglutinin and Neuraminidase Proteins": Table S8_PDBSUM_TLR7_MZA.docx

| Table S8. Detailed interaction summary for the vaccine–TLR7 complex | | |
| --- | --- | --- |
| Interaction  Type | Number of  Interactions | Interacting Residues Pairs (Chain A__ Chain B) |
| Salt Bridges | 2 | Lys383(chainA) __Glu279(chain B)  Lys110(chain A) __Glu297(chain B) |
| Disulphide Bonds | 0 | __ |
| Hydrogen Bonds | 16 | Tyr356 (Chain A) — Asn281 (Chain B) ,Tyr264 (Chain A) — Arg301 (Chain B) , Gln354 (Chain A) — Thr277 (Chain B) , Gln354 (Chain A) — Gln302 (Chain B) , Asn276 (Chain A) — Gly304 (Chain B) , Lys274 (Chain A) — Pro307 (Chain B) , Cys273 (Chain A) — Gly306 (Chain B) ,Pro272 (Chain A) — Pro305 (Chain B) ,Asn215 (Chain A) — Cys331 (Chain B) , Ser192 (Chain A) — Lys345 (Chain B) ,Thr271 (Chain A) — Gly334 (Chain B) , Arg194 (Chain A) — Gln344 (Chain B) ,Arg186 (Chain A) — Ser400 (Chain B) ,Asn187 (Chain A) — Cys401 (Chain B) ,Gln138 (Chain A) — Leu405 (Chain B) |
| Non-bonded Contacts | 213 | They are present throughout the docked complex. |
| This table summarizes the inter-chain interactions between Chain A and Chain B of the vaccine–TLR7 complex, as predicted by the PDBsum server. Number of interactions are shown in second column, and the interacting residues are shown in the third column. | | |
