## Supplementary Tables for "Immunoinformatics-Guided Design and In Silico Evaluation of a Multi-Epitope Vaccine Against Influenza A H10N5 and H3N2 Strains Based on Hemagglutinin and Neuraminidase Proteins": Table S9_Prodigy Results_TLR3_MZA.docx

**Table S9: Predicted binding affinity and interaction profile for the vaccine–TLR3 complex**

| Feature | Value | Interpretation |
| --- | --- | --- |
| ΔG (Binding Free Energy) | -16.1 kcal/mol | Very strong binding (lower is better; < -10 is considered strong) |
| Kd (Dissociation Constant) | 1.4 × 10⁻¹² M | Extremely tight binding (picomolar range means high affinity) |
| Interfacial Contacts (ICs) |  |  |
| Charged–Charged | 8 | Moderate salt bridge/electrostatic involvement |
| Charged–Polar | 14 | N/A |
| Charged–Apolar | 33 | N/A |
| Polar–Polar | 7 | N/A |
| Polar–Apolar | 34 | N/A |
| Apolar–Apolar | 34 | High hydrophobic interactions |
| NIS (Non-Interacting Surface) |  |  |
| Charged | 20.73% | N/A |
| Apolar | 39.27% | Significant hydrophobic contribution |
| This table shows the results of PROGIDY analysis for the vaccine complex bounded with the TLR3 complex as calculated by the PRODIGY server. Interaction features are listed in the first column, whereas the values and interpretations for each interactive feature are listed in second and third column. | | |
