## Supplementary Tables for "Immunoinformatics-Guided Design and In Silico Evaluation of a Multi-Epitope Vaccine Against Influenza A H10N5 and H3N2 Strains Based on Hemagglutinin and Neuraminidase Proteins": Table S10_Prodigy_MZA_TLR7.docx

| Table S11. Predicted Binding Affinity and interaction profile for the vaccine and TLR7 complex | | |
| --- | --- | --- |
| Feature | **Value** | **Interpretation** |
| ∆G (Binding Free Energy) | -16.8 | Very strong binding (lower is better;<-10 is considered stronger) |
| Kd (Dissociation Constant) | 4.6e-13 | Extremely tight binding  (Picomolar range means  High affinity) |
| Interfacial Contacts (ICs) |  |  |
| Charged-Charged | 21 | Moderate salt bridge  /Electrostatic involvement |
| Charged-Polar | 39 | N/A |
| Charged-Apolar | 55 | N/A |
| Polar-Polar | 12 | N/A |
| Polar-Apolar | 22 | N/A |
| Apolar-Apolar | 22 | High hydrophobic Interactions |
| NIS (non-Interacting surface) |  |  |
| Charged | 26.75 | N/A |
| Apolar | 29.75 | Significant Hydrophobic Interactions |
| This table shows the results of PROGIDY analysis for the vaccine complex bounded with the TLR7 complex as calculated by the PRODIGY server. Interaction features are listed in the first column, whereas the values and interpretations for each interactive feature are listed in second and third column. | | |
